# Septins coordinate cell wall integrity and lipid metabolism in a sphingolipid-dependent process

**DOI:** 10.1101/2020.02.09.940718

**Authors:** Alexander Mela, Michelle Momany

**Affiliations:** Fungal Biology Group and Plant Biology Department, University of Georgia, 2502 Miller Plant Science Building, Athens, GA 30602

**Keywords:** **S**eptins, MAPK Signaling, Cell wall integrity, Sterol Rich Domains

## Abstract

During normal development and response to environmental stress, fungi must coordinate synthesis of the cell wall and plasma membrane. Septins, small cytoskeletal GTPases, colocalize with membrane sterol-rich regions and facilitate recruitment of cell wall synthases during dynamic wall remodeling. In this study we show that null mutants missing an *Aspergillus nidulans* core septin present in hexamers and octamers (Δ*aspA^cdc11^*, Δ*aspB^cdc3^*, or Δ*aspC^cdc12^*) are sensitive to multiple cell wall-disturbing agents known to activate the cell wall integrity MAPK pathway and that this sensitivity can be remediated by osmotic support. The null mutant missing the octamer-exclusive core septin (Δ*aspD^cdc10^*) showed similar osmotic-remedial sensitivity, but only to a single cell wall-disturbing agent and the null mutant missing the noncore septin (Δ*aspE*) showed very mild osmotic-remedial sensitivity to a different single agent. Representative core septin null mutants showed changes in cell wall polysaccharide composition, organization, and chitin synthase localization. Double mutant analysis with *ΔmpkA* suggested core septins interact with the cell wall integrity pathway. Null mutants missing any of the five septins were resistant to ergosterol-disrupting agents. The Δ*aspA^cdc11^*, Δ*aspB^cdc3^*, and Δ*aspC^cdc12^* mutants showed increased sensitivity to sphingolipid-disrupting agents that was remediated by addition of exogenous phytosphingosine. Representative core septins were mislocalized after treatment with sphingolipid-disrupting agents, but not after treatment with ergosterol-disrupting agents. When challenged with both sphingolipid-disturbing and cell wall-disturbing agents in combination, remediation of the lipid defect restored proper growth to Δ*aspA^cdc11^*, Δ*aspB^cdc3^*, and Δ*aspC^cdc12^*, but remediation of the cell wall defect did not. Our data suggest that the core hexamer and octamer septins are involved in cell wall integrity signaling with the noncore septin playing a minor role; that all five septins are involved in monitoring ergosterol metabolism; that the hexamer septins are required for sphingolipid metabolism; and that septins require sphingolipids to coordinate the cell wall integrity response.

## Introduction

The cell wall and plasma membrane (PM) are the primary lines of defense against environmental insults for fungal cells. With the large surface area of hyphal networks and intimate contact with the surrounding media, fungi encounter many stresses, ranging from ion imbalance to predation. The cell wall contains several polysaccharide constituents: chitin provides the rigid framework of the cell wall, β-glucan maintains the shape, and mannans form the outermost, protective layer (1–6). Precise regulation of cell wall and plasma membrane architecture is attained by tightly coordinated signaling pathways which control genes responsible for maintaining homeostasis between the intracellular and extracellular environments.

Septins are highly conserved small GTPase cytoskeletal proteins that function as molecular scaffolds for dynamic cell wall and plasma membrane-remodeling, as well as diffusion barriers restricting movement of membrane and cell wall-associated molecules (7–14). The *Saccharomyces cerevisiae* septins Cdc3, Cdc10, Cdc11, and Cdc12 have been termed ‘core septins’ because they are monomers which assemble into non-polar heterooligomers and micrometer-scale higher order structures in the form of bars, rings, or gauzes (15–18). *A. nidulans* contains four orthologous septin proteins: AspA^Cdc11^, AspB^Cdc3^, AspC^Cdc12^, and AspD^Cdc10^, as well as AspE, which has no *S. cerevisiae* orthologue (19). *A. nidulans* contains two distinct subpopulations of heterooligomers; an octameric oligomer consisting of all four core septins in the same order as proposed in *S. cerevisiae* (AspA^Cdc11^-AspC^Cdc12^-AspB^Cdc3^-AspD^Cdc10^-AspD^Cdc10^-AspB^Cdc3^-AspC^Cdc12^-AspA^Cdc11^), and a second hexameric oligomer, with the proposed order (AspA^Cdc11^-AspC^Cdc12^-AspB^Cdc3^-AspB^Cdc3^-AspC^Cdc12^-AspA^Cdc11^)(12, 13, 20). For clarity, we will refer to AspA^Cdc11^, AspB^Cdc3^, and AspC^Cdc12^ as ‘core hexamer septins’; AspA^Cdc11^, AspB^Cdc3^, AspC^Cdc12^, and AspD^Cdc10^ as ‘core octamer septins’; and AspE as the ‘noncore septin’ because it does not assemble into oligomeric structures, though it is required for higher order structure assembly at the multicellular stage (20, 21).

Previous studies in *Candida albicans* have shown that septins provide the scaffolding for cell wall proteins via the **C**ell **W**all **I**ntegrity (CWI) MAPK signaling pathway (7, 9, 22–24). The cell wall integrity pathway, along with the other major MAPK signaling pathways (**H**igh **O**smolarity **G**lycerol (HOG), cAMP-PKA, **T**arget **o**f **R**apamycin (TOR), Calcineurin/Calcium, and Mating/Pheromone response pathways) are highly conserved across the Kingdom Fungi, and have been shown to undergo extensive cross-talk to coordinate virtually all biological functions in the cell, from expansion and division to asexual reproduction (25–32).

Sphingolipids are long chain base-containing lipids (33) that are metabolized in a highly conserved pathway in plants, animals, and fungi; sphingolipid metabolism shares direct connections to other major metabolic pathways, such as sterol metabolism and fatty acid and phospholipid synthesis (34). Sphingolipid pathway intermediates, such as phytosphingosine, have been shown to be involved in CWI pathway signaling in *S. cerevisiae* (33, 35). **S**terol **r**ich **d**omains (SRDs), also called ‘lipid rafts’ or ‘lipid microdomains,’ are regions of the plasma membrane enriched in specific classes of lipids, including sterols (ergosterol, the major sterol found in most fungi (36)), sphingolipids, and phosphoinositides (37, 38), which have been shown to be functionally important for maintenance of cell polarity (39). Membrane organization, plasticity, and overall integrity have been attributed to sphingolipid, sterol, and glycerophospholipid interactions (40). *In vitro* work has shown that yeast septins bind directly to the phosphoinositides PIP2, PI(4,5)P2, PI(5)P, and PI(4)P (41–44). Septins have also been shown to genetically interact with Sur2, a sphinganine C4-hydroxylase which catalyzes the conversion of sphinganine to phytosphingosine in sphingolipid biosynthesis (45), and to physically interact with GTPases, Bud1 and Cdc42 to maintain sphingolipid-dependent diffusion barriers at cell membranes in yeast (46). Interdependent colocalization of septins and sterol rich microdomains have been described in a number of *in vivo* and *in vitro* systems (47, 48). In the dimorphic fungus *U. maydis,* septins and sterol rich domains depend on each other to localize properly at the hyphal tip for cell signaling and cell polarity establishment (49). In the budding yeast *S. cerevisiae,* long-chain sphingolipids have been implicated in maintaining asymmetric distribution and mobility of some membrane-spanning molecules, such as multidrug resistance transporters (50). In contrast to these findings, a more recent comprehensive study of protein segregation during cell division in budding yeast found that the majority of ER and PM proteins with transmembrane domains were actually symmetrically segregated, however septins were shown to be responsible for partitioning of the PM-associated ER at the bud neck, thereby restricting diffusion of ER lumen and a particular set of membrane-localized proteins (51).

Recent work has started to unravel the functional connections between the septins, cell wall integrity MAPK pathway signaling, and lipid metabolism, however most studies have focused on a small sub-set of septin monomers and/or were conducted in primarily yeast-type fungi (52–57). Here we show in the filamentous fungus *A. nidulans* that the core septins, AspA^Cdc11^, AspB^Cdc3^, AspC^Cdc12^, and AspD^Cdc10^ are required for proper coordination of the cell wall integrity pathway, that all five septins are involved in lipid metabolism, and that these roles require sphingolipids.

## Results

### Mutants missing core hexamer septins are hypersensitive to multiple cell wall-disturbing agents. The mutant missing the octamer-exclusive core septin AspD^Cdc10^ is sensitive to a single agent and the mutant missing the noncore septin AspE is mildly sensitive to a single agent

To determine whether *A. nidulans* septins are important for cell wall integrity, we used spore dilution assays to test the growth of septin deletion mutants on media containing a variety of known cell wall-disturbing agents. Wild type and septin null mutants were tested on Calcofluor White (CFW) and Congo Red (CR), cell wall polymer-intercalating agents that perturb chitin and β-1,3-glucan, respectively; Caspofungin (CASP), an inhibitor of β-1,3-glucan synthase; and Fludioxonil (FLU), a phenylpyrrol fungicide that antagonizes the group III histidine kinase in the osmosensing pathway and consequently affects cell wall integrity pathway signaling (**Fig 1**)(58–67). Septin mutants Δ*aspA^cdc11^*, Δ*aspB^cdc3^*, and Δ*aspC^cdc12^* showed hypersensitivity to CFW, CR, CASP, and FLU; Δ*aspD^cdc10^* displayed no sensitivity to CFW, CR, and FLU, and sensitivity to CASP (See **S1 Table** for list of strains used in this study). The Δ*aspE* mutant showed no sensitivity to CFW, CASP, and FLU, and very mild sensitivity to CR. (**Fig 1A**).

**Fig 1.**
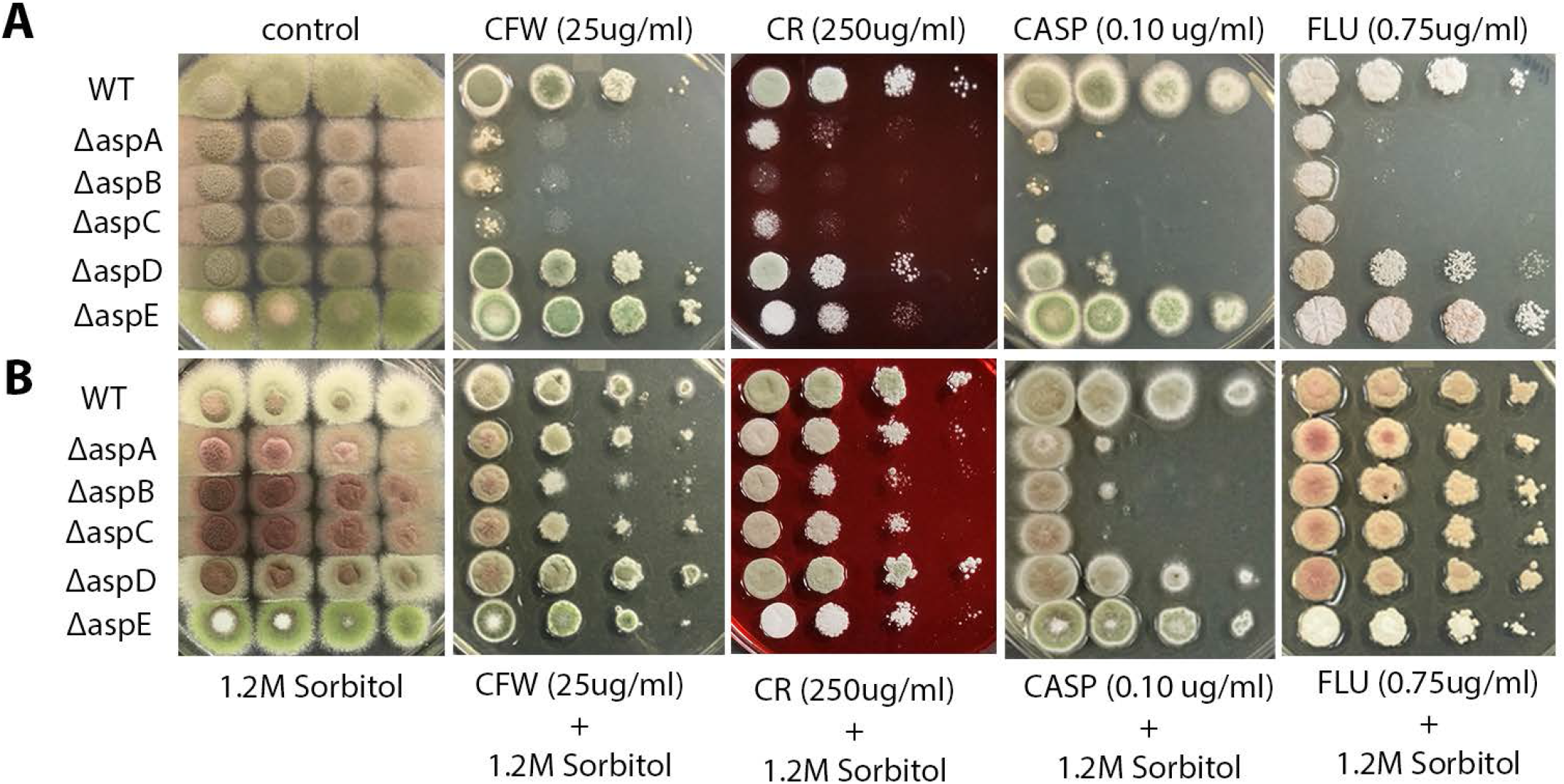
Septin null mutants exhibit sensitivity to cell wall-disturbing agents, which can be remediated by osmotic support. (Top Row) Solid media spotting assay. WT and septin null mutants Δ*aspA^cdc11^*, Δ*aspB^cdc3^*, Δ*aspC^cdc12^*, Δ*aspD^cdc10^*, and Δ*aspE* were tested for sensitivity by spotting decreasing spore concentrations on complete medium plates with or without cell wall-disturbing agents Calcofluor White (CFW), Congo Red (CR), Caspofungin (CASP), and Fludioxonil (FLU) at the indicated final concentrations. (Bottom Row) WT and septin null mutants were tested for osmotic remediation of hypersensitivity to cell wall-disturbing agents, by spotting decreasing spore concentrations on media amended with exogenous 1.2M sorbitol. Spore concentrations were 10^7^ conidia/mL – 10^4^ conidia/mL for all assays. Differences in colony color result from changes in spore production, spore pigment, and production of secondary metabolites under stress. N≥5

### Septin mutant sensitivity to cell wall-disturbing agents is remediated by osmotic stabilization

One hallmark of cell wall integrity defects is rescue by the addition of an osmotic stabilizer, such as sorbitol or sucrose. The addition of exogenous 1.2M sorbitol partially remediated the hypersensitivity to CASP and fully remediated the hypersensitivity to CFW, CR, and FLU for Δ*aspA^cdc11^*, Δ*aspB^cdc3^*, and Δ*aspC^cdc12^* (**Fig 1B**). Sorbitol also remediated the sensitivity of Δ*aspD^cdc10^* to CASP and the mild sensitivity of Δ*aspE* to CR. The osmotic remediation of growth defects in the mutants indicates that their sensitivity to cell wall-disturbing agents is likely the result of a cell wall integrity defect.

### Core septin null mutant Δ*aspB^cdc3^* has altered chitin and β-1,3-glucan localization

Because the core septin mutants showed sensitivity to cell wall-disturbing agents consistent with action via the cell wall integrity pathway, we predicted that there might be differences in cell wall polymer localization in the mutants. To examine cell wall polymer localization, we did live-cell imaging of WT and Δ*aspB^cdc3^* since it is a member of both hexamers and octamers. Conidia were incubated on coverslips in liquid media, stained with CFW or aniline blue to observe chitin and β-1,3-glucan, respectively, and immediately observed by fluorescence microscopy. Z-stack images were analyzed one-by-one, compressed into maximum projection images to visualize the fluorescence signal in the entire hyphal structure, and by line scans of aniline blue and calcofluor white staining patterns (**Fig 2, S1 Fig**). As previously reported, Δ*aspB^cdc3^* showed more presumptive branch initials per hyphal compartment than WT (21). The aniline blue staining showed a reduction of labeling at all Δ*aspB^cdc3^* hyphal tips. Smaller, less intense aniline blue labeling also occurred at presumptive branch initials (**Fig 2A, S1A**). The CFW staining of Δ*aspB^cdc3^* showed a shift of the tip band of staining closer to the hyphal apex and a less well-defined endocytic collar zone (**Fig 2B, S1B**). These data show that cell wall organization is altered in Δ*aspB^cdc3^* and raise the possibility that it might be altered in other core hexamer septin null mutants as well.

**Fig 2.**
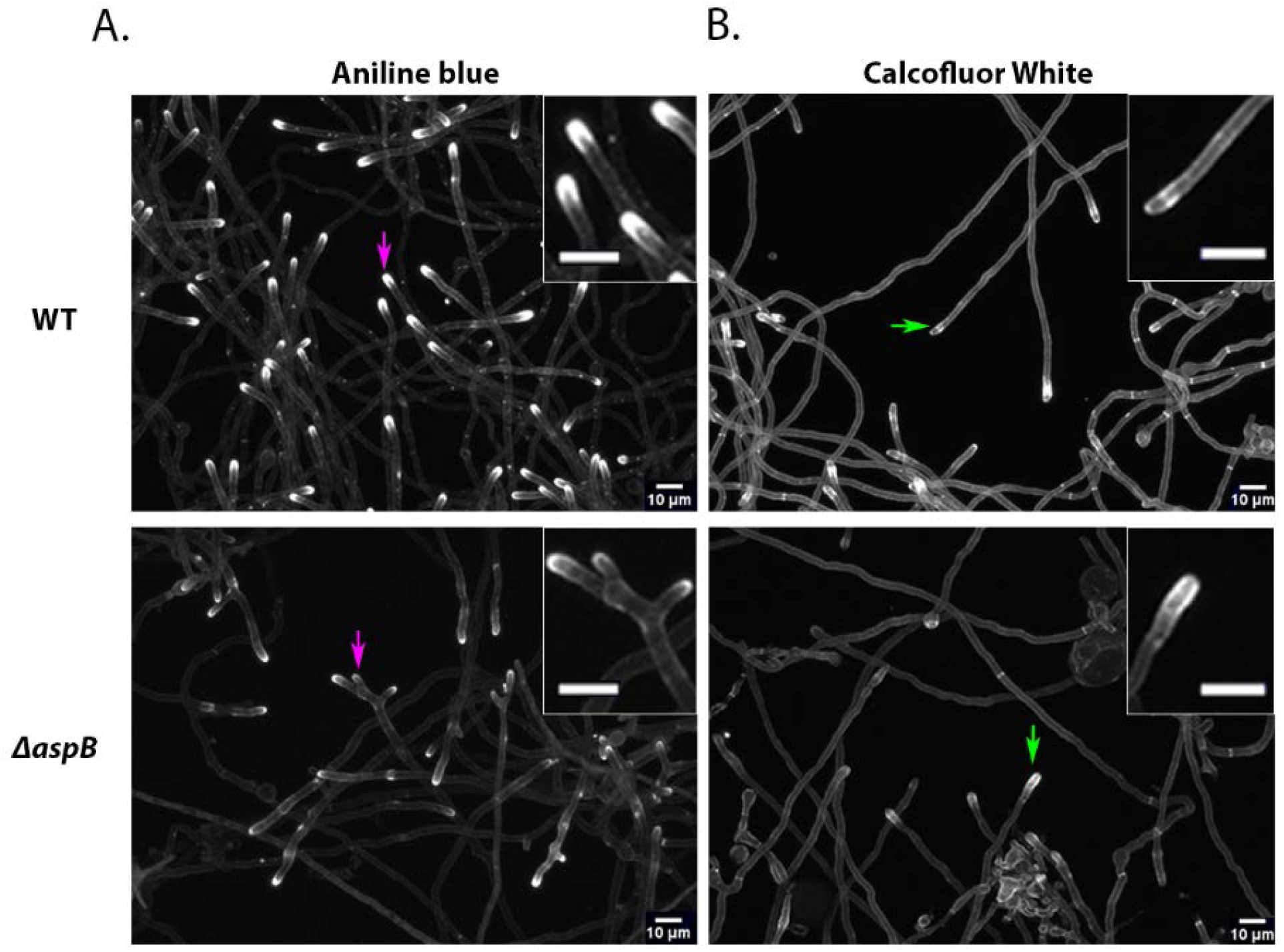
Cell wall staining reveals unique patterns of chitin and β-1,3-glucan deposition in the core septin hexamer null mutant Δ*aspB^cdc3^*. (A) WT and Δ*aspB^cdc3^* cells were incubated for approximately 14h and (A) stained with aniline blue to visualize β-1,3-glucan or (B) stained with Calcofluor White (CFW) to visualize chitin. Representative images are shown from at least three independent biological replicates, with ≥100 cells observed each. Magenta arrows in panel A denote hyphal tips with representative β-1,3-glucan deposition. Green arrows in panel B denote hyphal tips with representative chitin deposition. Insets show enlargement of area around the arrows. Scale bars = 10μm. N≥3

### Core septin null mutant Δ*aspB^cdc3^* has higher levels of chitin

Previous studies have shown that perturbations to one cell wall component often trigger compensatory changes to others via the cell wall integrity pathway (28, 68, 69). To analyze the cell wall composition of septin mutants, two independent biological replicates of a diagnostic glycosyl linkage analysis (70) were conducted to quantify the cell wall polysaccharide composition of WT, Δ*aspB^cdc3^* (a core septin null mutant which showed hypersensitivity to cell wall-disturbing agents), and Δ*aspE* (non-core septin mutant)(**S2 Fig**). All samples contained 3- and 4-linked glucose as the primary cell wall components, as well as 4-linked N-acetylglucosamine (the monomer of chitin) and a relatively minor amount of mannan. The septin mutants Δ*aspB^cdc3^* and Δ*aspE* showed 7% and 2% increases in the average area of the detected linkage peak for 4-GlcNAc compared to WT, respectively; the amount of 4-Glc and of mannan (data not shown) did not show significant differences between samples. These data indicate an increase in chitin content of the Δ*aspB^cdc3^* mutant and raise the possibility that it might be altered in other core hexamer septin null mutants as well.

### Chitin synthase localization is altered in the core septin null mutant ΔaspA^cdc11^

Membrane-spanning cell wall synthases are the ultimate effectors of the cell wall integrity pathway. *A. nidulans* contains six genes for chitin synthases: *chsA, chsB, chsC, chsD, csmA*, and *csmB*. Chitin synthase B localizes to sites of polarized growth in hyphal tips, as well as developing septa in vegetative hyphae and conidiophores, a pattern very similar to septin localization. Deletion of chitin synthase B shows severe defects in most filamentous fungi analyzed thus far, and repression of the chitin synthase B gene expression in *chsA, chsC*, and *chsD* double mutants exacerbated growth defects from a number of developmental states observed in each single mutant, suggesting it plays a major role in chitin synthesis at most growth stages (22). For these reasons, we chose chitin synthase B as a candidate to observe in a septin mutant background for possible defects in localization. We hypothesized that recruitment or maintenance of cell wall synthases, such as chitin synthase could be disrupted in septin null mutants which showed sensitivity to cell wall-disturbing agents and altered cell wall staining patterns. To determine the localization of synthases, a chitin synthase B-GFP (*chsB-GFP*) strain was crossed with strains in which core hexamer septins were deleted. After repeated attempts, the only successful cross was with core hexamer deletion strain Δ*aspA^cdc11^*. The Δ*aspA^cdc11^, chsB-GFP* strain showed conspicuous differences in chitin synthase localization patterns compared to WT (**Fig 3 and Fig. S3**). In WT, the chitin synthase-GFP signal was at the tips of >90% of branches, presumptive branch initials, and apical hyphal tips (S Fig 3A). In the septin null mutant Δ*aspA^cdc11^*, GFP signal was absent in most presumptive branch initials, but present in longer side branches that were at least 10 μm long and in the apical hyphal tip(s) (**Fig 3 and Fig. S3A**). Intriguingly, the chitin label calcofluor white was more intense in the septin deletion strain consistent with the increase in 4-GlcNAc levels seen in polysaccharide analysis (Fig. S3B). Calcofluor white labeling also showed that the polymer chitin was present throughout the hyphae in both the WT and septin deletion strains (Fig. 2) suggesting that chitin synthases had been active in both, but perhaps were not retained as long in the presumptive branch initials of the septin deletion mutant. These data support the hypothesis that core hexamer septins play a role in recruitment and/or retention of cell wall synthases to the proper locations at the plasma membrane during growth and development.

**Fig 3.**
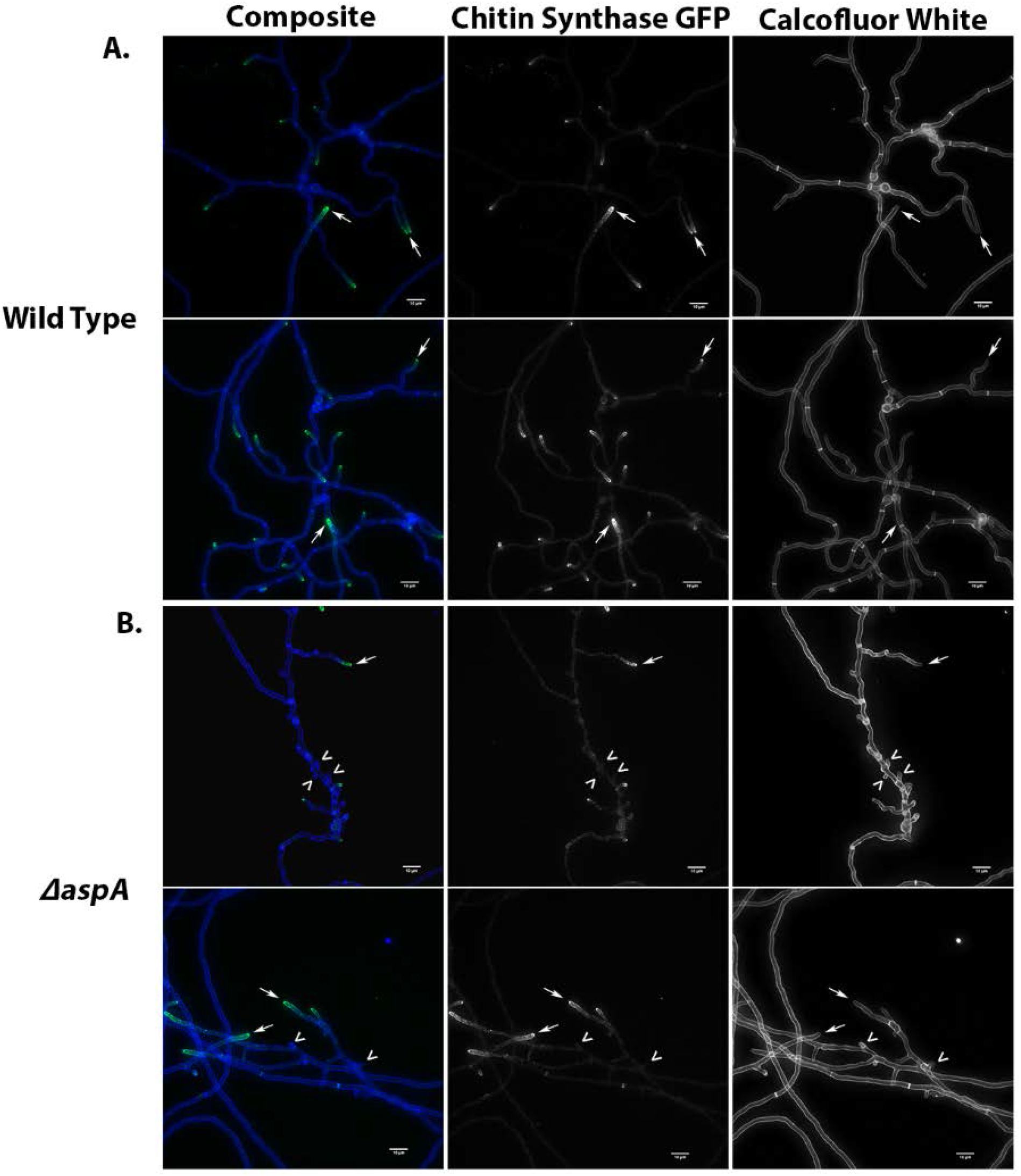
Chitin synthase is mislocalized in core septin null mutant Δ*aspA^cdc11^*. (A) Chitin synthase B-GFP in WT background (Columns 1-2). (B) chitin synthase B-GFP in a Δ*aspA^cdc11^* mutant background. Calcofluor White labeling shows the presence of the polymer chitin at septa, main hyphal tips, branches, and putative branch initials (Column 3). Representative images are shown from at least three independent biological replicates, with ≥100 cells observed. White arrows highlight hyphal branches and tips. White arrow heads highlight putative branch initials. Scale bars = 10μm. N≥3.

### Double mutant analyses suggest the core septin aspB^cdc3^ modulates the cell wall integrity pathway in the ΔmpkA background under cell wall stress

To determine whether there are genetic interactions between the septins and the cell wall integrity pathway kinases, double mutants were generated by sexual crosses and analyzed. The first cell wall integrity pathway kinase, PkcA^Pkc1^ (ANID_00106), is essential in *A. nidulans,* and so null alleles could not be utilized for double mutant analysis. A null allele of the final kinase in the CWI MAPK signaling cascade, MpkA^Slt2^ (ANID_11786), was crossed with Δ*aspB^cdc3^* and Δ*aspE* and progeny were challenged with cell wall-disturbing treatments. If the septins are directly in the CWI pathway, we would expect double mutants to show the same phenotypes as the parental null mutant that acts earliest. If the septins are in a parallel pathway or alternate node which also affects cell wall integrity, we would expect a novel/synergistic phenotype in the double mutants.

Spore dilution assays were conducted, challenging the double mutants and the parental strains with cell wall-disturbing treatments. The double mutants Δ*aspB^cdc3^* Δ*mpkA^slt2^* and Δ*aspE* Δ*mpkA^slt2^* displayed a colony-level radial growth defect and reduced conidiation which phenocopied the Δ*mpkA^slt2^* single mutant. When challenged with low concentrations of CASP and CFW, the Δ*aspB^cdc3^* Δ*mpkA^slt2^* and Δ*aspE* Δ*mpkA^slt2^* mutants were more sensitive than Δ*aspB^cdc3^* and Δ*aspE* single mutants, but suppressed the colony growth defects of Δ*mpkA^slt2^*. The novel phenotype of the double mutants shows that septins are involved in cell wall integrity and raises the possibility that they act in a bypass or parallel node for remediation of cell wall defects (**Fig 4**).

**Fig 4.**
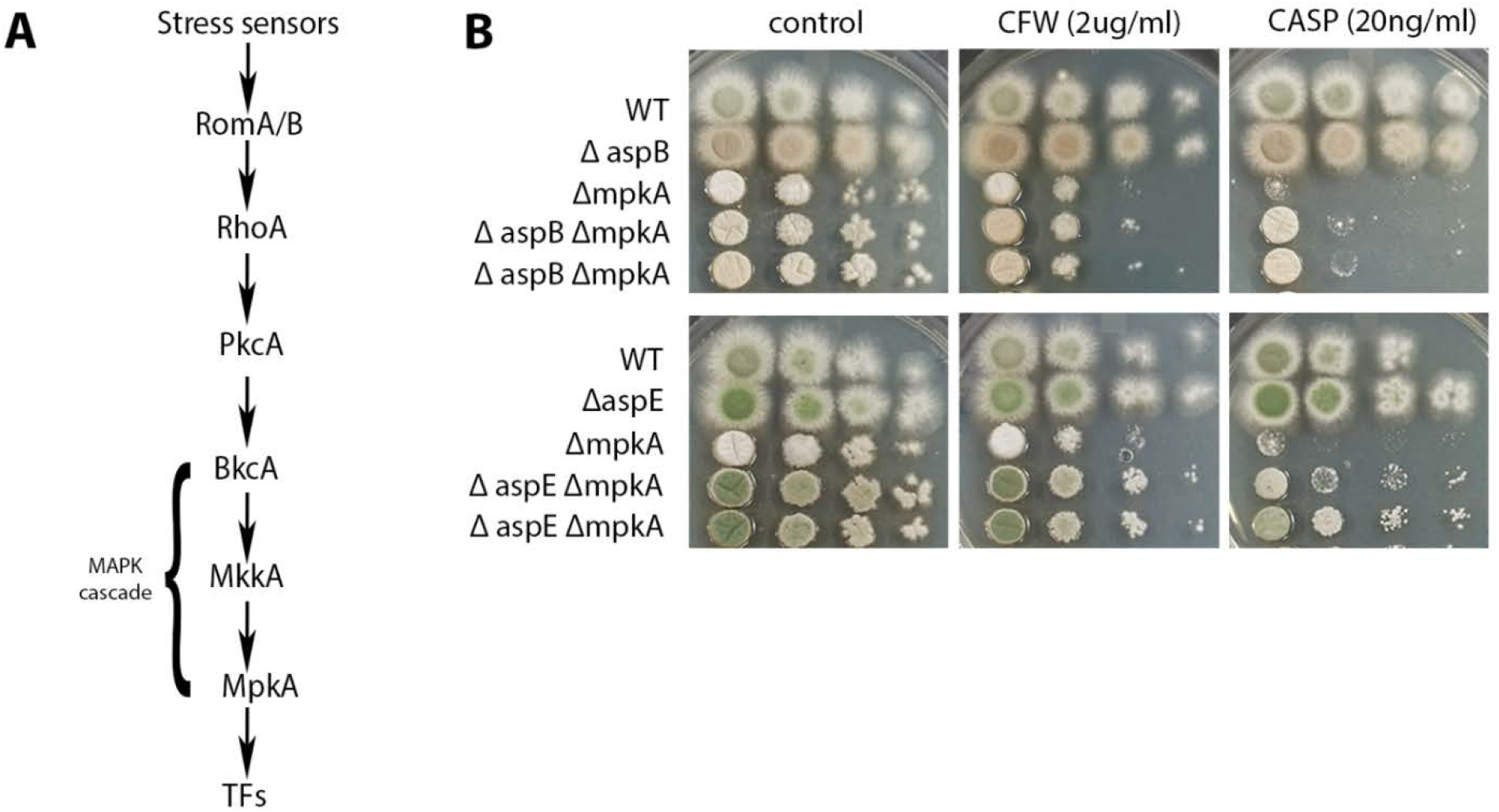
Double mutant analyses suggest core septins modulate the cell wall integrity pathway. (A) Simplified schematic diagram of the *A. nidulans* cell wall integrity MAPK signaling pathway (71). (B) Solid media spotting assays. (Top) WT, Δ*aspB^cdc3^*, Δ*mpkA^slt2^*, and two Δ*aspB^cdc3^* Δ*mpkA^slt2^* double mutant strains, were tested for sensitivity by spotting decreasing spore concentrations on complete media plates with or without cell wall-disturbing agents. (Bottom) WT, Δ*aspE*, Δ*mpkA^slt2^*, and two Δ*aspE* Δ*mpkA^slt2^* double mutant strains were tested for sensitivity by spotting decreasing spore concentrations on complete media plates with or without cell wall-disturbing agents. Differences in colony color result from changes in spore production, spore pigment, and production of secondary metabolites under stress. Transcription Factors, (TFs); Calcofluor White, (CFW); Caspofungin (CASP). Spore concentrations were [10^6^ conidia/mL – 10^3^ conidia/mL] for all assays in figure. N=3

### Core septin null mutants are insensitive to treatments which disrupt the Ca2+/Calcineurin, cAMP-PKA, or TOR Pathways. Noncore septin null mutant ΔaspE is sensitive to some TOR pathway inhibitors

One possible explanation for the observed sensitivity to cell wall-disturbing agents could be that septins are involved in ‘cross-talk’ with other MAPK signaling pathways that have been shown to interact with cell wall integrity pathway signaling such as the Calcium/Calcineurin (CAMK) signaling pathway. To test this possibility, calcium chloride, EGTA (calcium chelating agent), and FK-506 (calcineurin inhibitor) were added to each treatment (**S4 Fig**) (72–74). The treatments showed no obvious colony growth defects, suggesting that Ca2+/Calcineurin signaling pathway crosstalk does not significantly contribute to the observed sensitivity of septin null mutants to cell wall-disturbing agents.

Another pathway closely associated with cell wall integrity, lipid biosynthesis, and lipid signaling is the TOR MAPK signaling pathway (75). To test the involvement of septins in this signaling pathway, septin null mutants were challenged with rapamycin (a potent inhibitor of the TOR pathway), as well as methylxanthine derivatives and TOR pathway inhibitors, caffeine and theophylline (76–78). There were no observable growth defects in the presence of rapamycin (final concentration was 600ng/mL, which is approximately 3-fold higher than inhibitory concentration for known TOR pathway mutants) in the septin null mutants compared to WT (**S4 Fig**) (79, 80). Caffeine and theophylline have been shown to interfere with phosphodiesterase activity in the cAMP-PKA and TOR pathways, and Δ*pkaA^tpk1^* and Δ*tor 1^torA^* mutants show hypersensitivity to caffeine treatment which cannot be remediated by exogenous sorbitol (58, 78, 81–83). If the septins were involved in crosstalk between the cAMP-PKA or TOR pathways and the CWI pathway, we would predict that septin null mutants would be hypersensitive to caffeine and theophylline treatments. To our surprise, only Δ*aspE* showed hypersensitivity to both caffeine and theophylline, and the hypersensitivity was not remediated by an osmotic stabilizer (**S4 Fig and data not shown**). These data suggest that TOR, cAMP-PKA, and Calcium/Calcineurin signaling pathways do not contribute to the cell wall sensitivity or plasma membrane resistance phenotypes in the core septin null mutants, but the cAMP-PKA or TOR pathways may contribute to the phenotypes of the septin Δ*aspE* null mutant.

### Core septin null mutants show increased resistance to disruption of ergosterol biosynthesis and Δ*aspA^cdc11^* and ΔaspE show increased resistance to the ergosterol-binding drug Natamycin

We hypothesized that septins could indirectly modulate MAPK pathways, particularly the CWI pathway, through interactions with plasma membrane lipids within sterol rich domains (SRDs). Recent work has shown that the cell wall integrity pathway in *S. cerevisiae* is regulated by sphingolipids and ergosterol, facilitating proper deposition of cell wall polymers at actively growing regions and sites of septation (38, 55). It is well-established that septins localize to sites of polarized growth and septation, where highly dynamic remodeling of the plasma membrane and cell wall via membrane-bound synthases and polarity-associated proteins takes place (84). These highly dynamic plasma membrane SRDs in fungi often contain ergosterol and sphingolipids. To determine whether loss of septins might affect one of the major SRD-associated lipids, each septin mutant was challenged by drug treatments disrupting the ergosterol biosynthesis pathway in spore dilution assays (**Fig 5**). These assays included an azole treatment (Itraconazole), an allylamine treatment (Terbinafine), and a polyene treatment (Natamycin), which each impact a different step in ergosterol biosynthesis (**Fig 5A**). Allylamines (Terbinafine) inhibit the conversion of squalene to squalene epoxide, azoles (Itraconazole) inhibit the conversion of lanosterol to 4,4-dimethylcholesta-8,14,24-trienol, and polyenes bind directly to ergosterol (85).

**Fig 5.**
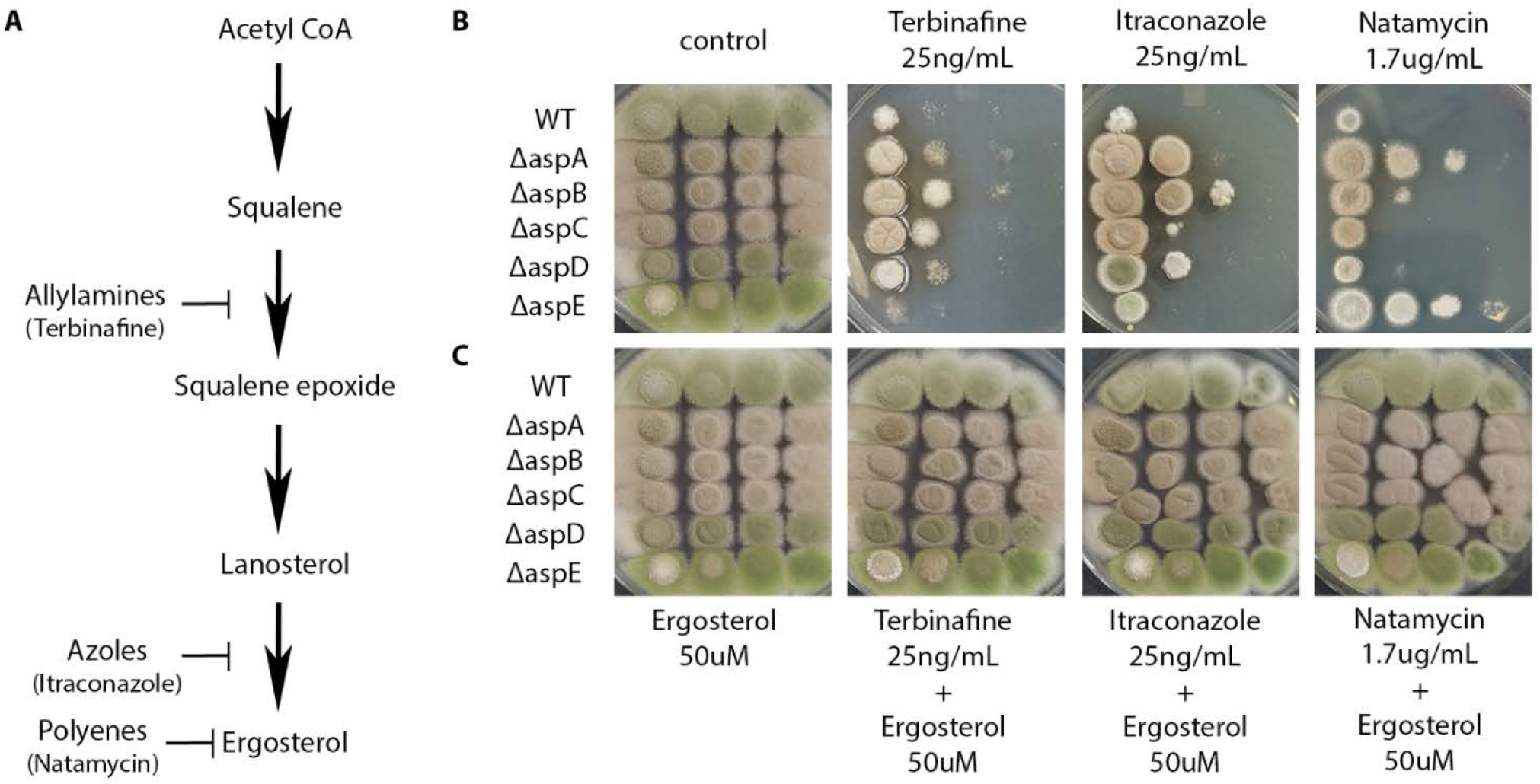
Core septin null mutants show increased resistance to disruption of ergosterol biosynthesis and Δ*aspA^cdc11^* and Δ*aspE* show increased resistance to the ergosterol-binding polyene drug Natamycin. (A) A simplified schematic diagram of the ergosterol biosynthesis pathway, showing where allylamines, azoles, and polyene antifungal agents affect each step (85). (B) WT and septin null mutants Δ*aspA^cdc11^*, Δ*aspB^cdc3^*, Δ*aspC^cdc12^*, Δ*aspD^cdc10^*, and Δ*aspE*, were tested for sensitivity by spotting decreasing spore concentrations on solid media with or without the ergosterol biosynthesis-disturbing agents Terbinafine, Itraconazole, and Natamycin at concentrations shown. (C) Remediation of sensitivity to ergosterol biosynthesis-inhibiting treatments. WT and septin null mutants were tested for the ability of exogenous ergosterol (50μM) to remediate sensitivity to ergosterol biosynthesis-disturbing agents. Differences in colony color result from changes in spore production, spore pigment, and production of secondary metabolites under stress. Spore concentrations were [10^7^ conidia/mL – 10^4^ conidia/mL] for all assays in figure. N=3

The Δ*aspA^cdc11^*, Δ*aspB^cdc3^*, Δ*aspC^cdc12^*, and Δ*aspD^cdc10^* mutants were more resistant to Itraconazole and Terbinafine than WT or Δ*aspE*. Only Δ*aspA^cdc11^* and Δ*aspE* showed strong resistance to Natamycin treatment (**Fig 5B**). The addition of exogenous 50μM ergosterol was able to fully remediate the sensitivity of all null mutant strains and WT to Itraconazole, Terbinafine, and Natamycin, suggesting ergosterol was indeed the primary lipid component disrupted by these treatments (**Fig 5C**). These data suggest that all five septins are involved in monitoring ergosterol metabolism and/or deposition.

### Core hexamer septin null mutants show altered sensitivity to disruption of sphingolipid biosynthesis

Sphingolipids are a class of plasma membrane lipids which has been shown to be associated with sterol-rich domains, along with sterols and phosphoinositides, and therefore are possible targets for septin-mediated interactions at the membrane. To determine whether loss of septins impacts sphingolipid metabolism in *A. nidulans*, the septin null mutants were challenged with sphingolipid biosynthesis-disrupting agents Myriocin and Aureobasidin A (AbA) (**Fig 6**). Myriocin disrupts the first committed step of the biosynthetic pathway, converting palmitoyl-coA and serine to 3-ketodihydrosphingosine, preventing the accumulation of downstream intermediates, such as ceramides and sphingoid bases like phytosphingosine (PHS), as well as complex sphingolipids at the final steps of the pathway (**Fig 6A**) (37, 47). Aureobasidin A inhibits IPC synthase, disrupting the conversion of inositolphosphorylceramide from phytoceramide, and consequently causing the accumulation of intermediates such as phytosphingosine, which has been shown to be toxic at high concentrations (**Fig 6A**) (86).

**Fig 6.**
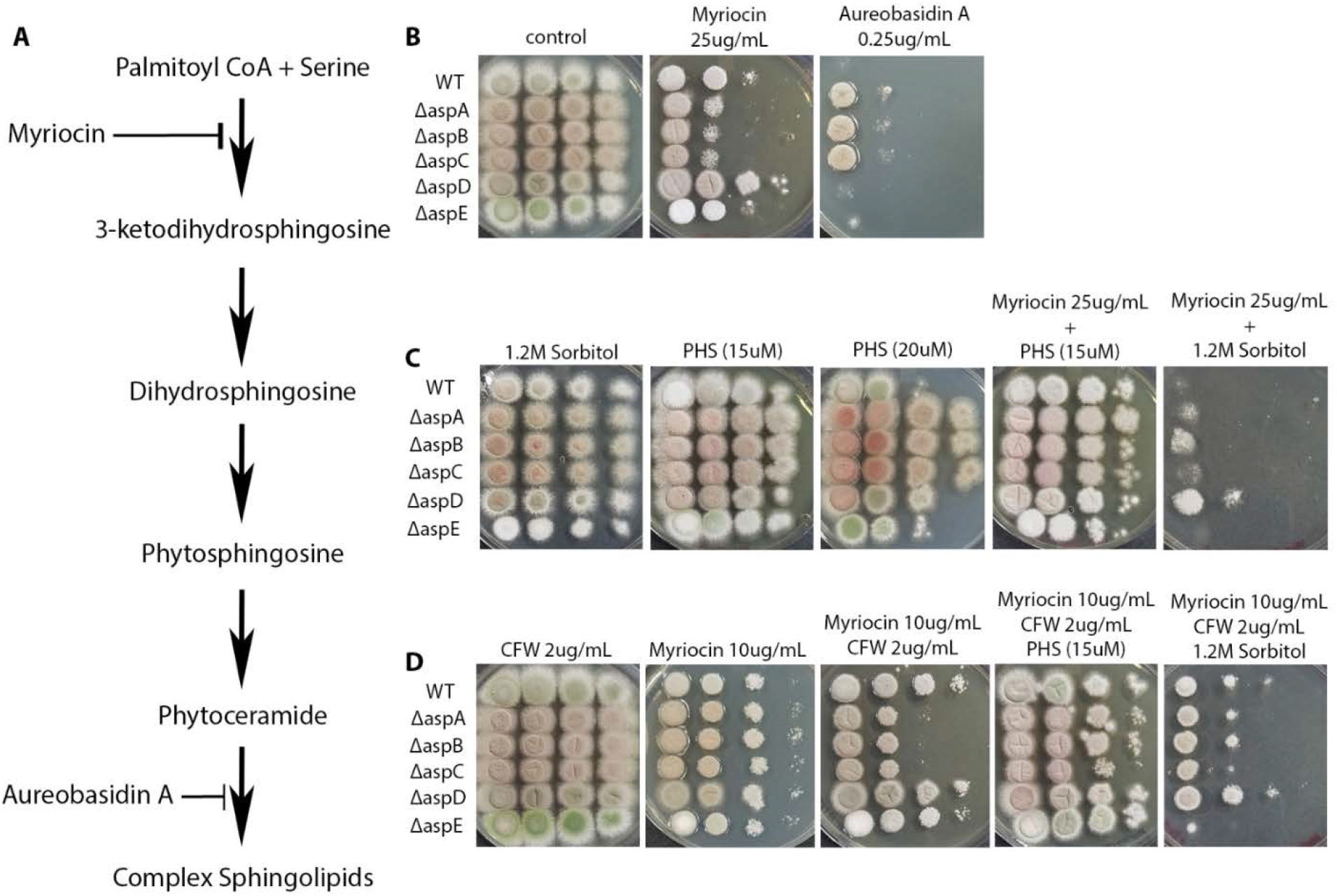
Septin core hexamer null mutants show altered sensitivity to agents which disrupt sphingolipid biosynthesis. (A) A simplified diagram of the sphingolipid biosynthesis pathway in *A. nidulans* (87). (B) *Solid media spotting assay*. WT and septin null mutants, Δ*aspA^cdc11^*, Δ*aspB^cdc3^*, Δ*aspC^cdc12^*, Δ*aspD^cdc10^*, and Δ*aspE* were tested for sensitivity by spotting decreasing spore concentrations on solid media with or without sphingolipid biosynthesis-disturbing agents Myriocin and Aureobasidin A. (C) *Remediation of sensitivity to sphingolipid biosynthesis-disturbing agents*. WT and septin null mutants were tested for remediation of sensitivity to sphingolipid biosynthesis-disturbing agents by spotting decreasing spore concentrations on solid media amended with exogenous phytosphingosine (PHS) intermediate (15μM) or 1.2M sorbitol, with or without Myriocin. (D) Combinatory treatment with cell wall and sphingolipid biosynthesis-disturbing agents. WT and septin null mutants were tested for sensitivity to cell wall and sphingolipid biosynthesis-disturbing agents in combination, by spotting decreasing spore concentrations on solid media with or without ‘sub-lethal’ concentrations of Calcofluor White (CFW), Myriocin, or CFW + Myriocin, amended with either exogenous phytosphingosine intermediate (15uM) or 1.2M sorbitol. Differences in colony color result from changes in spore production, spore pigment, and production of secondary metabolites under stress. Spore concentrations were [10^7^ conidia/mL – 10^4^ conidia/mL] for all assays in figure. N=3

The core hexamer null mutants Δ*aspA^cdc11^*, Δ*aspB^cdc3^*, and Δ*aspC^cdc12^* were more sensitive to Myriocin than the other septin null mutants or WT (**Fig 6B**). Δ*aspA^cdc11^*, Δ*aspB^cdc3^*, and Δ*aspC^cdc12^* were also more resistant to AbA than null mutants of the octamer-exclusive septin Δ*aspD^cdc10^*, the noncore septin Δ*aspE*, or WT (**Fig 6B**). Strikingly, the addition of exogenous PHS (15μM) to the Myriocin treatment fully remediated the sensitivity of Δ*aspA^cdc11^*, Δ*aspB^cdc3^*, and Δ*aspC^cdc12^* (**Fig 6C**). Δ*aspA^cdc11^*, Δ*aspB^cdc3^*, and Δ*aspC^cdc12^* were also more resistant to higher concentrations (20μM) of the phytosphingosine intermediate (PHS), which has been shown to be toxic at high concentrations (**Fig 6C**).

To address whether cell wall and plasma membrane defects might be associated with one another in septin null mutants, a combinatory drug treatment approach was taken. Sublethal concentrations of CFW (2μg/mL) and Myriocin (10μg/mL), in which all strains grew at every spore concentration, were combined (**Fig 6D**). When combined, the two drugs resulted in additive, colony-level growth defects for Δ*aspA^cdc11^*, Δ*aspB^cdc3^*, and Δ*aspC^cdc12^*. This sensitivity to both drugs in combination was remediated by the addition of exogenous PHS. Surprisingly the addition of exogenous sorbitol, which had fully remediated the hypersensitivity of the septin null mutants to all previously tested cell wall-disturbing agents, resulted in a more dramatic growth defect in media containing only Myriocin, or in combination with CFW (**Fig 6C and D**). These data together suggest that the core hexamer septins AspA^Cdc11^, AspB^Cdc3^, and AspC^Cdc12^ may monitor sphingolipid metabolism. They further suggest that the hexamer septins may signal sphingolipid status to the cell wall integrity pathway, and that this signal is required for proper cell wall integrity function.

### Localization of core hexamer septins AspA^Cdc11^ and AspB^Cdc3^ is disrupted by sphingolipid inhibitors

We predicted that since core septin hexamer null mutants Δ*aspA^cdc11^*, Δ*aspB^cdc3^*, and Δ*aspC^cdc12^* were sensitive to drugs which inhibit sphingolipid biosynthesis the localization of these septins might be altered when exposed to the same treatments. We examined whether disruption of sphingolipid biosynthesis causes changes in septin localization using live-cell imaging with fluorescence microscopy. A strain carrying AspA^Cdc11^-GFP was grown in liquid complete medium overnight, treated with exogenous PHS (15μM), and imaged for 3 hours post-treatment. There was a dynamic shift in septin localization under PHS treatment over the course of the experiment, compared to the vehicle control. The septin-GFP signal shifted from a relatively homogenous cortical localization along the hyphal tips to a more stochastic, punctate localization along the entire length of hyphae (**Fig 7, middle panel**).

**Fig 7.**
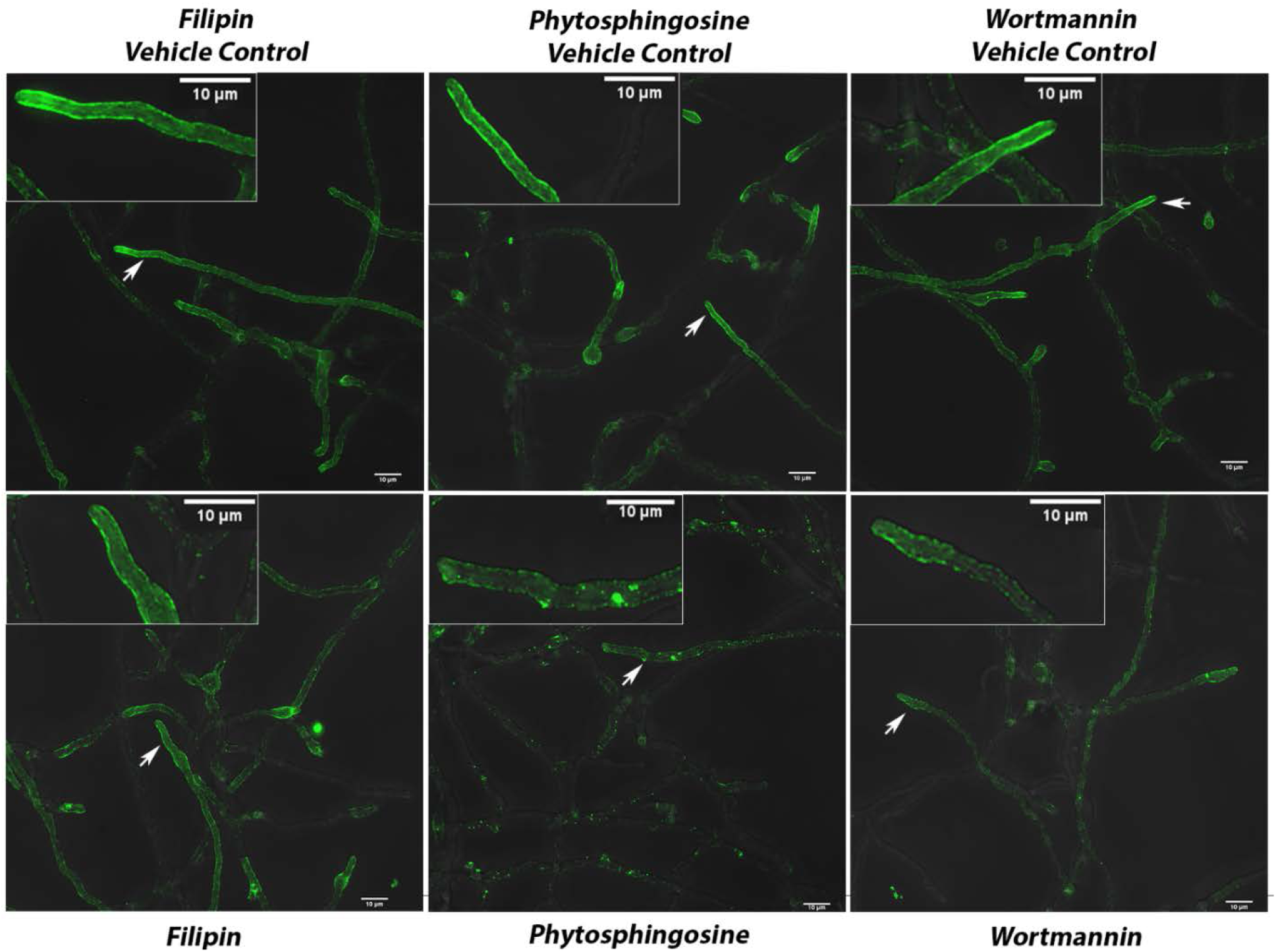
Septin AspA^Cdc11^-GFP localization disrupted by sphingolipid biosynthesis inhibitors. AspA^Cdc11^-GFP strain incubated in liquid media for approximately 16h and imaged 180 minutes after replacing with fresh media containing sterol, sphingolipid, and phosphoinositide-disturbing agents, Filipin, phytosphingosine, and Wortmannin, respectively. Representative images are shown from three independent biological replicates with ≥100 cells observed. (Top Row) Vehicle controls for each treatment. (Bottom Row) Filipin (25μg/mL), phytosphingosine (15μM), and Wortmannin (20μg/mL) treatments respectively. Insets show enlarged section of micrographs from each picture to better visualize pattern of fluorescence signal. Imaging conducted with Deltavision I deconvolution inverted fluorescence microscope. White arrows denote hyphae which are highlighted in enlarged images. Scale bars = 10μm. N=3

In contrast to these results, treatments with the ergosterol-binding polyene, Filipin III (25μg/mL) and phosphatidylinositol 3-kinase inhibitor, Wortmannin (20μg/mL), did not affect the localization pattern of septins as dramatically as the PHS treatment (**Fig 7, left- and right-most panels**). Similar patterns of aberrant localization were observed under PHS treatments for AspB^Cdc3^-GFP (**S5 Fig**). Myriocin treatment also resulted in a similar pattern of mislocalization in AspA^Cdc11^-GFP cells when compared to treatment with PHS (**S6 Fig**). These results suggest sphingolipid content and/or distribution within the plasma membrane contributes to the localization and stability of core hexamer septins at the plasma membrane and that ergosterol and phosphoinositides may not be as vital for this process.

## Discussion

Our data show that *A. nidulans* septins play roles in both plasma membrane and cell wall integrity and that distinct subgroups of septins carry out these roles. Previous work has shown that the five septins of *A. nidulans* septins form hexamers (AspA^Cdc11^, AspB^Cdc3^, and AspC^Cdc12^) and octamers (AspA^Cdc11^, AspB^Cdc3^, AspC^Cdc12^, and AspD^Cdc10^) and that the noncore septin AspE does not appear to be a stable member of a heteropolymer (20). The current work suggests that though all septins are involved in coordinating cell wall and membrane integrity, the roles of hexamers, octamers, and the noncore septin are somewhat different. Core hexamer septins appear to be most important for sphingolipid metabolism, all five septins appear to be involved in ergosterol metabolism, and core septins are most important for cell wall integrity pathway with the noncore septin possibly playing a minor role. As summarized in Figure 8 and discussed in more detail below, our previous and current data are consistent with a model in which: (A) All five septins assemble at sites of membrane and cell wall remodeling in a sphingolipiddependent process; (B) All five septins recruit and/or scaffold ergosterol and the core hexamer septins recruit and/or scaffold sphingolipids and associated sensors at these sites, triggering changes in lipid metabolism; and (C) The core septins recruit and/or scaffold cell wall integrity machinery to the proper locations and trigger changes in cell wall synthesis. The noncore septin might play a minor role in this process.

**Fig 8.**
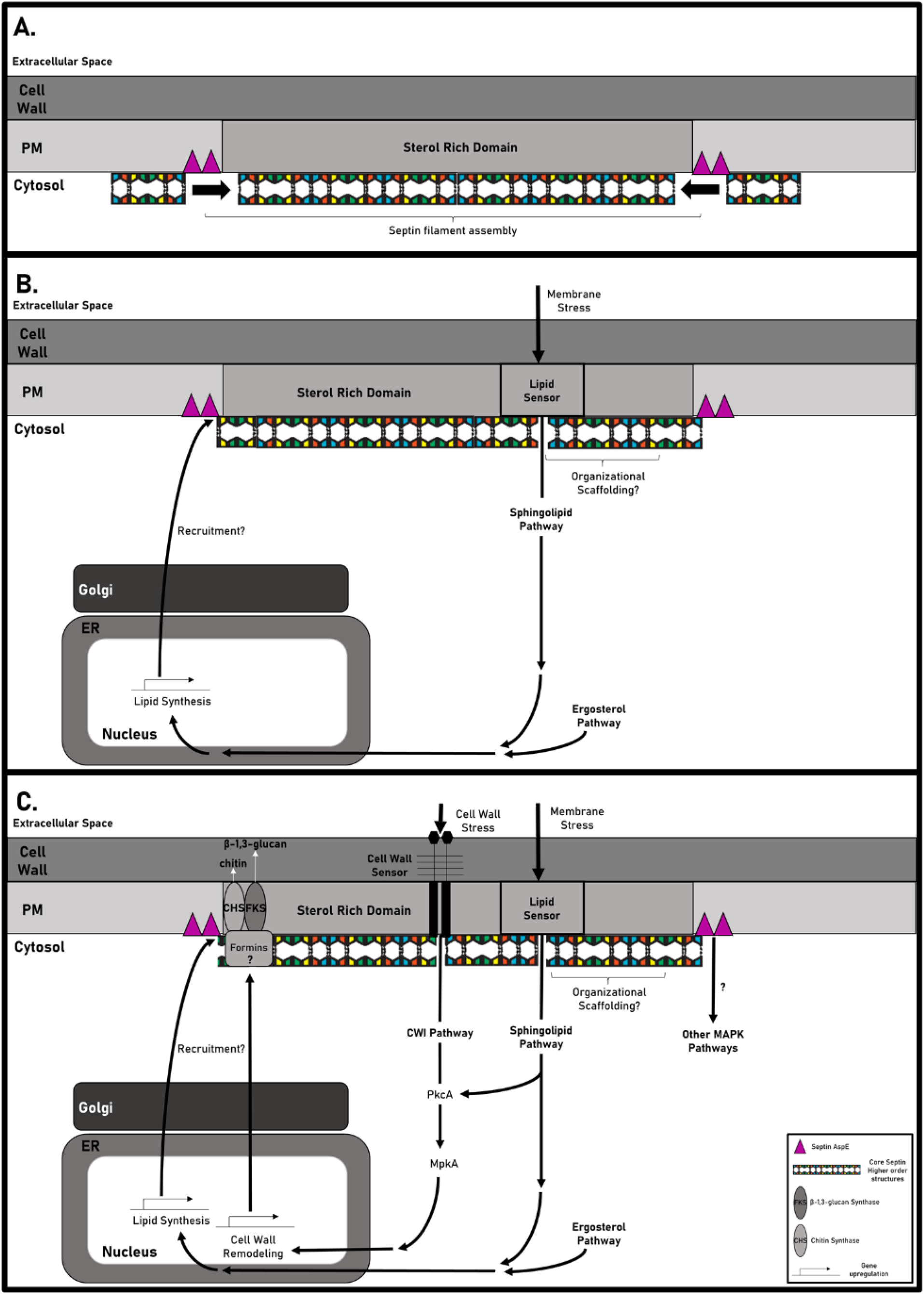
Model for septin modulation of cell wall and plasma membrane integrity through interactions with sterol rich domains (SRDs). As described in the text, our data suggest that all five septins are involved in cell wall and membrane integrity coordination. The core septins that participate in hexamers appear to be most important for sphingolipid metabolism while all septins appear to be involved in ergosterol metabolism and cell wall integrity. Because SRDs contain both sphingolipids and ergosterol and because it is not yet clear how subgroups of septins interact with each other at SRDs, we show all core septins in our model without distinguishing hexamers and octamers. In this model, septins are proposed to colocalize with SRDs in a manner which promotes: (A) assembly of septins into higher order structures along the membrane at sites of polarized growth or cell wall/PM remodeling; (B) the recruitment and/or scaffolding of lipids and associated membrane-bound sensors to monitor membrane composition and/or stress. The status of membrane composition is relayed to the nucleus where it triggers changes in expression of genes responsible for lipid metabolism; and (C) the recruitment and/or scaffolding of cell wall integrity pathway machinery to monitor cell wall composition and stress, followed by the recruitment and/or scaffolding of cell wall synthases, possibly with the help of other septin-interacting proteins such as formins. Question marks (?) denote speculative processes or interactions which have not been characterized.

### Septins assemble at sites of membrane and cell wall remodeling in a sphingolipid-dependent process

We hypothesize that sterol rich domain-associated lipids (ergosterol, sphingolipids, and phosphoinositides) recruit or facilitate binding and assembly of septins on the membrane, with sphingolipids contributing more to the stabilization of septins than other SRD-associated lipids in *A. nidulans* (**Fig 8A**). Previous studies showed preferential *in vitro* binding of yeast core septin orthologues to the phosphoinositides PIP2, PI(4,5)P2, PI(5)P, and PI(4)P, however our treatments with Wortmannin and Filipin III, known disruptors of phosphoinositides and ergosterol respectively (88–92), did not affect septin localization as dramatically as sphingolipid-disturbing treatments (**Fig 7, S3, and S4 Fig**). The marked septin mislocalization we observed upon treatment with phytosphingosine and Myriocin strongly supports the idea that sphingolipids at the membrane contribute to core septin localization and help maintain septin assemblies at the proper locations. Previous studies on composition of lipid microdomains showed that relatively minor differences in sphingolipid structure can have significant effects on sterol and phospholipid interactions, consequently resulting in major changes in membrane properties (93, 94). Perhaps sphingolipids (and ergosterol to a lesser extent) help to stabilize SRDs in a way that facilitates assembly of septin filaments and higher order structures via diffusion, collision, and annealing as proposed by Bridges et al (2014) (41–43, 95). Consistent with this idea, *A. nidulans* AspB filaments have been shown to move along the plasma membrane, break apart, and ‘snap’ together in a way that suggests collision and annealing (21).

### Septins recruit and/or scaffold lipids and associated sensors, triggering changes in lipid metabolism

Consistent with a role for septins in modulating membrane composition, all septin null mutants were resistant to ergosterol-disrupting treatments and core hexamer septin mutants were affected by disruption of sphingolipid biosynthesis (**Fig 5A-B, Fig 6**). Based on proposed mechanisms for septins as diffusion barriers or organizational scaffolds of membrane-associated proteins in yeast and smut fungi, we propose septins monitor lipid microdomain composition and/or organization in filamentous ascomycetes (49, 51)(**Fig 8B**). Septins do not have a transmembrane domain, a feature that often defines established membrane ‘sensors’ that monitor local lipid environments (96, 97); however, septins share a highly conserved polybasic domain proposed to facilitate septin-membrane interactions (44, 98). In addition to the polybasic domain, septins have recently been shown to contain an amphipathic helix motif which has been implicated in septin sensing of membrane curvatures (99). Given that septins have been shown to assemble into non-polar higher order structures along the plasma membrane (95), septin assembly itself might be the mechanism by which lipid composition and protein organization is monitored at the cytosolic face of membranes. Perhaps septin assemblies that pass specific size or geometric thresholds trigger signaling through MAPK and other pathways.

### Core septins recruit and/or scaffold cell wall integrity machinery to the proper locations and trigger changes in cell wall synthesis

In addition to monitoring and relaying information about the membrane, septins clearly have a role in normal cell wall growth and remediation of cell wall stress via the cell wall integrity pathway. We propose a major role of the core septins is to recruit and/or organize integral proteins to sites of polarized growth or remodeling at the cell cortex to ensure cell wall integrity pathway functions are carried out (**Fig 8C**). The sensitivity to cell wall-disturbing agents, altered cell wall composition, and altered polysaccharide deposition in the core septin null mutants (**Fig 1–2** and **S1–S2 Fig**) are consistent with phenotypes of cell wall integrity pathway mutants in previous studies (100–102).

Our glycosyl linkage analysis showed that cell wall chitin content is increased in the septin mutants compared to WT (**S2 Fig**). Hyper-synthesis of chitin has been shown to occur during cell wall stress via the cell wall integrity pathway in *S. cerevisiae* and *Candida albicans* (103–110). Double mutant analyses between septins and CWI pathway kinases also support a role for core septins in maintaining cell wall integrity under stress (**Fig 4**). Suppression of cell wall defects under cell wall stress by deletion of septins in an Δ*mpkA^slt2^* background suggests a parallel node by which septins negatively regulate cell wall integrity pathway sensors or kinases could exist. This interpretation is consistent with studies in yeast showing Bni4 (ANID_00979), a formin which is phosphorylated by and functions downstream of MAPK *Slt2^mpkA^*, directly interacts with the core septin orthologues in order to recruit chitin synthases to the bud neck (111–113). Though the involvement of AspE in the cell wall integrity pathway appears to be minor based on the very mild sensitivity of the null mutant to Congo Red (**Fig 1**), sensitivity to TOR and cAMP-PKA pathway inhibitors suggests that AspE might participate indirectly through MAPK pathway crosstalk (**Fig S4**). It could be that in the septin null mutants, septin filaments are not able to properly assemble and monitor lipid composition and protein organization at the cytosolic face of membranes, resulting in defects in plasma membrane and cell wall integrity (Fig 8).

Though we have discussed membrane and cell wall integrity separately, it is possible that that membrane defects in the septin null mutants contribute to the observed cell wall changes or that cell wall defects contribute to the observed changes in lipid metabolism. When septin deletion mutants were challenged with both membrane-disturbing and cell wall-disturbing agents in combination, remediation of the lipid defect (via PHS) restored proper growth, but remediation of the cell wall defect (via sorbitol) did not remediate lethality (**Fig 6C-D**). This suggests that there is a synergistic effect of disrupting sphingolipid metabolism and cell wall architecture in septin null mutants and that septin-sphingolipid interactions are required for roles in maintaining cell wall integrity.

Because *A. nidulans* septin deletion mutants are viable, we were able to systematically analyze the roles of all septins in this organism. Our data show that all septins are required for proper coordination of lipid metabolism, with core hexamer septins most important for sphingolipid metabolism and all septins involved in ergosterol metabolism. Our data also show that the core septins are most important for cell wall integrity pathway, but that these roles require sphingolipids. Based on our data we propose that septins are critical for tight coordination of plasma membrane metabolism and cell wall synthesis during normal development and response to exogenous stress and that the site of this coordination is likely SRDs. Future work will address how subgroups of septins interact with each other and with sphingolipids and ergosterol in SRDs to coordinate the cell wall integrity response.

## Methods

### Spore Dilution Drug Sensitivity Assays

Strains used in this study are listed in **S1 Table**. Media used were previously described (114). *Aspergillus nidulans* strains were harvested in ddH20, spores normalized to 1X10^7^ or 1X10^6^ conidia/mL, and then serial diluted 4-fold into separate Eppendorf tubes. All strains were inoculated in 10uL droplets in a grid pattern on petri plates containing 25mL of solid (1.8% agar) complete medium (CM) (1% glucose, 0.2% peptone, 0.1% Casamino Acids, 0.1% yeast extract, trace elements, nitrate salts, and 0.01% vitamins, pH 6.5; with amino acid supplements as noted) or minimal medium (MM)(1% glucose, trace elements, 1% thiamine,0.05% biotin, pH 6.5; with amino acid supplements as noted) with or without amended supplements. All incubations were conducted at 30°C as indicated for 3-4 days before images were taken. Sorbitol, NaCl, and KCl were added at 2M, 1M, and 1.5M respectively to media before autoclaving. Stock solutions were prepared as follows: Blankophor BBH/Calcofluor White (Bayer Corporation; standard-SV-2460; 25mg/mL in ddH20 and adjusted pH with 1M KOH until solubilized), Congo Red (Fisher Scientific; Lot No.8232-6; 10mg/mL in ddH20), Caspofungin acetate (1mg/mL in ddH20), Fludioxonil/Pestanal (2mg/mL in DMSO), and caffeine monohydrate (63.66mg/mL in ddH20 and gently heated with stirring until solubilized); calcium chloride (2M in ddH20), EGTA (0.5M in ddH20), Rapamycin/Sirolimus (1mg/mL in acetonitrile), FK-506 (5mg/mL in DMSO), Natamycin (1.5mg/mL in MeOH), Itraconazole (10mg/mL in DMSO), Terbinafine (10mg/mL in DMSO). Stock solutions of Myriocin (5mg/mL), phytosphingosine (1mg/mL), and Aureobasidin A (1mg/mL) were prepared in DMSO, EtOH, and MeOH respectively and stored at −20C in the dark. Images of plates were captured using a cellular device with an 8.0 Megapixel camera, and subsequently processed using Photoshop CS5 Version 12.0 X32.

### Generating double mutants and other strains by crossing

Parental strains were coinoculated in CM liquid at 1X10^5^ conidia/mL supplemented with all parental auxotrophic markers and allowed to incubate at 30°C for up to one week or until a thick mycelial mat had formed. Mycelial mats were transferred to solid MM plates containing only shared auxotrophic markers from the genetic backgrounds of each parental strain. Plates containing mycelial mats were parafilmed and incubated in the dark at room temperature for up to 2 weeks or until mature cleistothecia form on the mycelial mats. Multiple cleistothecia from each genetic cross were collected in water, diluted, and plated onto solid media containing all auxotrophic supplements from each parental strain to allow growth of all resulting progeny. Approximately 50 progenies were collected from each dissected cleistothecium, and each colony was transferred to master plates, and replica-plated onto minimal media without any supplements, to isolate prototrophic progeny. Five to ten progenies from each cross were then 3-phased streaked to obtain single colony isolates for PCR verification. All progenies of genetic crosses were verified by diagnostic PCR using KOD XTREME Hot Start DNA Polymerase (71975-3, EMD Millipore) or OneTaq^®^ Hot Start Quick-Load^®^ 2X Master Mix with Standard Buffer (M0488L, New England BioLabs inc.) according to manufacturer’s instructions. Double mutant strains of *ΔmpkA^slt2^* were verified for deletion of *mpkA^slt2^* by amplification of entire gene using primers, MpkA-806-F’ and MpkA-3779-R’, followed by SacI HF restriction enzyme digestion of PCR product to better visualize band sizes. All progeny from chitin synthase and septin null mutant crosses were determined to be virtually identical to each septin mutant parental strain in growth/morphology on a colony level and by microscopy, PCR-verified for the deletion of each septin gene, and screened visually by fluorescence microscopy for the presence of chitin synthase-GFP signal. Chitin synthase-GFP strains in a *ΔaspA^cdc11^* genetic background were verified for deletion of *aspA^cdc11^* by amplification of the entire gene using primers, AspA-KO-F’ and AspA-KO-R’, followed by XhoI restriction enzyme digestion for verification of band sizes. Strains and primer sets used in this study can be found in S1 and S2 Tables respectively.

### Growth conditions and microscopy

Growth and preparation of cells were as previously reported (115). Conidia were inoculated on sterile coverslips in 10mL liquid complete or minimal media at 1X10^5^ conidia/mL and incubated at 30C in a small petri dish for the specified amount of time. Cell walls were stained for chitin with Blankophor BBH (CFW) (American Cyanamid, Wayne, NJ; 25mg/mL stock solution in ddH2O and pH adjusted by 1M KOH until solubilized; working solution made by diluting stock solution by 100X and 8ul dissolved in 5mL ddH20 prepared fresh for working solution and used immediately), β-(1,3)-glucans were stained with aniline blue (stock solution prepared fresh to 10 mg/mL final concentration in ddH20; working solution prepared at 135.55μM in 50mM phosphate buffer, pH adjusted to 9.5 with 5M KOH, and used immediately; coverslips stained for 5 minutes in the dark at RT). Live cell imaging experiments tracking septin localization were conducted using Filipin III (stock solution prepared at 5mg/mL in DMSO; working solution was used at 25μg/mL final concentration in liquid complete media), phytosphingosine (working solution was used at 15μM in liquid complete media), and Wortmannin (stock solution prepared to 2mg/mL in DMSO; working solution was used at 20ug/mL in liquid complete media). Vehicle controls and conducted at ≤1% w/v in liquid media. Imaging was performed in the Biomedical Microscopy Core at the University of Georgia.

Microscopy was carried out using Zeiss Axioplan microscope and Zeiss Axiocam MRc charge-coupled device camera and software, as well as Deltavision I Deconvolution Inverted Microscope and LSM880 Confocal Fluorescent microscope with Diode laser (405nm), Argon (458, 488, 514nm) and HeNe (543, 633nm) laser lines. All micrograph comparisons between treatments imaged with identical microscope settings. Subsequent image analysis and scale bars added to micrographs, using ImageJ software 1.48v, Java 1.6.0_20 (64-bit) or Zen 2.3 imaging software, and final figures compiled in Photoshop CS5 software version 12.0 x32.

### Quantification of aniline blue and CFW staining patterns by line scans

Max projection images and scale bars were made in each micrograph in Zen (Black) Edition. Micrographs were false colored black and white, line scans were drawn by hand through hyphal tips, line scan profiles were generated in Zen 3.1 (Blue) imaging software, and associated date points for each profile were exported to Microsoft Excel version 2004 (Build 12730.20270) to create graphs and conduct statistical analysis. Figures were generated using Adobe Photoshop 2020 Version: 21.1.2.

### Quantification of chitin synthase-GFP and CFW fluorescence signal by line scans

Max projection images were made for each micrograph, line scans were drawn by hand through hyphal branch tips in ImageJ software 1.48v, and data points were exported to Microsoft Excel version 2004 (Build 12730.20270) to create graphs and conduct statistical analysis. Figures were assembled using Adobe Photoshop 2020 Version: 21.1.2.

### Cell Wall Extraction

Cell walls were isolated from a protocol based on (Bull, 1970); cell wall extraction and lyophilization were conducted as previously described in (Guest and Momany, 2000) with slight modifications listed in full procedure here. A single batch of complete media (recipe described above) was autoclaved and supplements added: Arginine, Methionine, Pyridoxine, and Riboflavin, were added to single flask and then distributed to individual flasks to be inoculated. 1 × 10^4^ conidia/mL were inoculated in 2 flasks each of 100mL liquid Complete Media. Flasks were incubated at 30C in orbital shaker at 200rpm for 48 hours. Mycelia was gravity filtrated, then vacuum-filtrated through #42 Whatman filter paper, subsequently washed with 50mL each of chilled ddH20 to remove residual media and stored at −80C until completely frozen. Mycelium from each sample was allowed to thaw on ice and then washed sequentially with 50mL chilled ddH20 and 0.5M NaCl. Fungal hyphal mats were transferred to mortar and pestle, then subsequently flash frozen in liquid nitrogen and ground in chilled Tris/EDTA Disruption Buffer (DB; 20mM Tris, 50mM EDTA, pH 8.0) with pre-chilled mortar and pestle. Samples were monitored by microscopy under 60X or 100X objective until hyphal ghosts were evident. Cell walls were separated by centrifugation at 13,800g for 10 min at 4C. Cell pellet was placed in a beaker with 40-100mL of chilled Tris/EDTA buffer and stirred at 4C for 12 hours. Cell pellet was collected by centrifugation as above and stirred again with 100mL chilled ddH20 at 4C for 4 hours. Cell wall materials was collected by vacuum filtration, frozen at −80C, lyophilized to dryness, and stored at room temperature (25°C) for further analysis.

### Cell Wall Glycosyl Linkage Analysis

To determine the glycosyl linkages, the samples were acetylated using pyridine and acetic anhydride in order to get better solubility, before two rounds of permethylation using sodium hydroxide (15 min) and methyl iodide (45 min). The permethylated material was hydrolyzed using 2M TFA (2 h in sealed tube at 121°C), reduced with NaBD4, and acetylated using acetic anhydride/pyridine. The resulting PMAAs (Neutral sugars) were analyzed on an Agilent 7890A GC interfaced to a 5975C MSD (mass selective detector, electron impact ionization mode); separation was performed on a 30 m Supelco SP-2331 bonded phase fused silica capillary column using Supelco SP-2331 fused silica capillary column (30 m x 0,25 mm ID). The PMAAs of amino sugars were separated on Supelco Equity-1 column (30 m x 0.25 mm ID). Further, the relative quantities of respective glycosyl linkages were calculated by integrating the peak area of respective peak. Since the neutral and amino sugars were analyzed on different instruments, the peak area of amino sugars was normalized with 4-Glc peak, which is prominent in both the instruments and the integrated and normalized peak areas were pooled together to calculate the relative percentage of individual linkages. Two independent, biological replicates were conducted for this analysis and processed in tandem. The average area (%) of detected linkages of one representative data set is included in the graphs to show relative differences between cell wall polysaccharides between samples.

## Acknowledgements

These studies were supported by the University of Georgia Franklin College of Arts and Sciences support to MM and Plant Biology Department support to AM. Chitin synthase strains were generously provided by Dr. Hiroyuki Horiuchi in the Department of Biotechnology at the University of Tokyo. The glycosyl linkage analysis was supported by a Chemical Sciences, Geosciences and Biosciences Division, Office of Basic Energy Sciences, U.S. Department of Energy grant (DE-SC0015662) to Parastoo Azadi at the Complex Carbohydrate Research Center at UGA.

## Supporting information

**S1 Fig.**
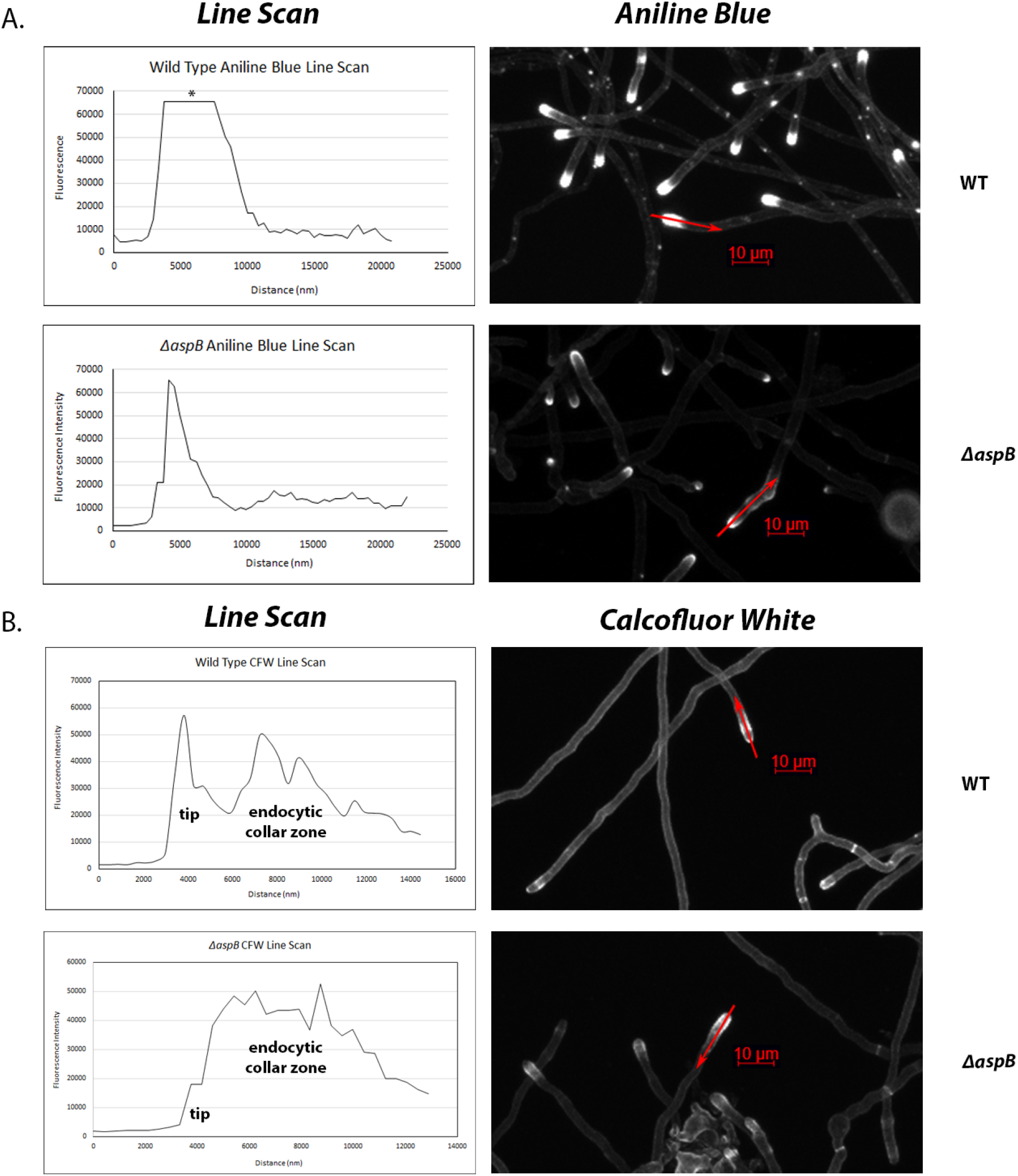
Line scans of cell wall staining reveals unique patterns of chitin and β-1,3-glucan deposition in core septin null mutants. (A) Line scan profiles (Left column) of aniline blue staining patterns, corresponding to the red arrows drawn through the hyphal tips of WT and Δ*aspB^cdc3^* (Right column). (B) Line scan profiles (Left column) of calcofluor white staining patterns, corresponding to the red arrows drawn through the hyphal tips of WT and Δ*aspB^cdc3^* (Right column). Asterisk in Panel A line scan profile denotes saturation of fluorescence signal.

**S2 Fig.**
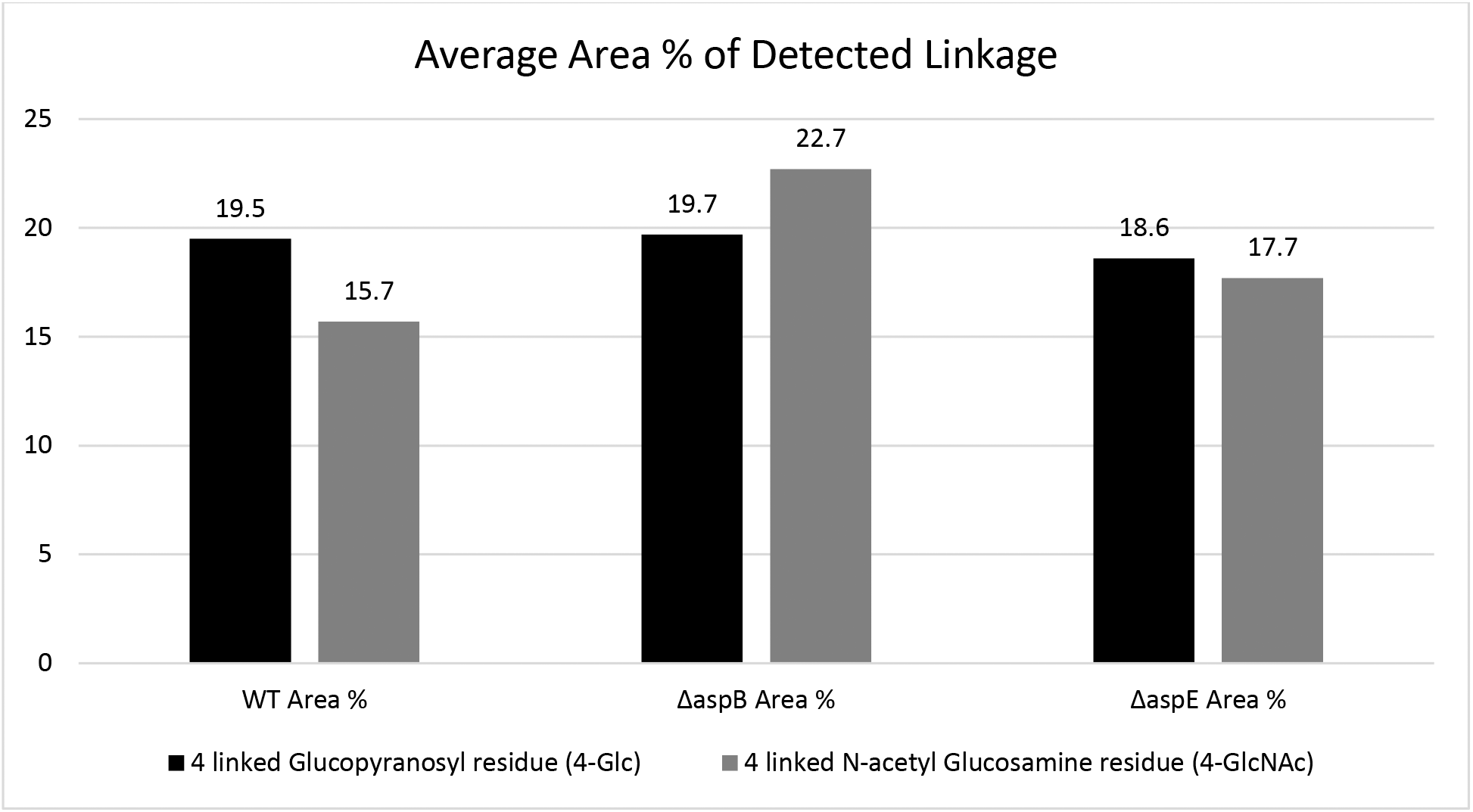
Cell wall glycosyl linkage analysis shows increased chitin content in Δ*aspB^cdc3^* septin null mutant. Results of cell wall polysaccharide glycosyl linkage analysis using GC MS/MS showing the average area (%) of detected linkages of 4-linked glucose and 4-linked N-acetyl glucosamine. Two independent biological replicates gave similar results. A representative data set is shown. Samples: WT, Δ*aspB^cdc3^*, and Δ*aspE*.

**S3 Fig.**
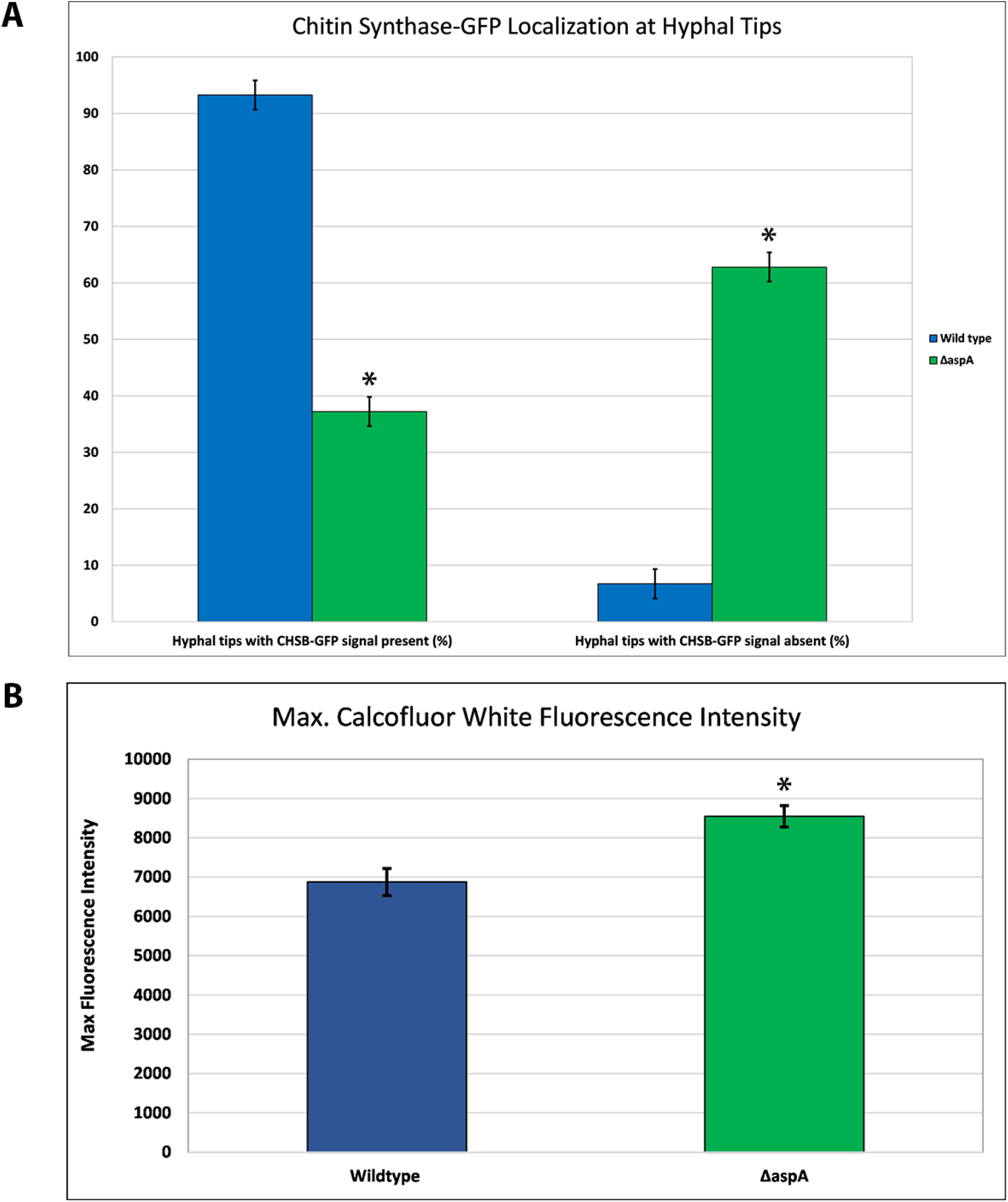
Quantification of chitin synthase B-GFP and CFW fluorescence signal. (A) Quantification of percentage (%) of hyphal branch tips with chitin synthase B-GFP present vs. absent by line scans in WT and Δ*aspA^cdc11^*. Mean values shown for line scans of ?85 hyphal branches over two independent biological replicates. Asterisks indicate different standard error of the mean between sample sets. (B) Quantification of maximum fluorescence intensity by line scans of CFW stained WT and Δ*aspA^cdc11^* hyphae (subapical region). Mean values shown for line scans of ≥65 hyphal branches over two independent biological replicates. Asterisks indicate different standard error of the mean between sample sets and statistically different values by two-tailed Ttest (p < 0.05) (N=50).

**S4 Fig.**
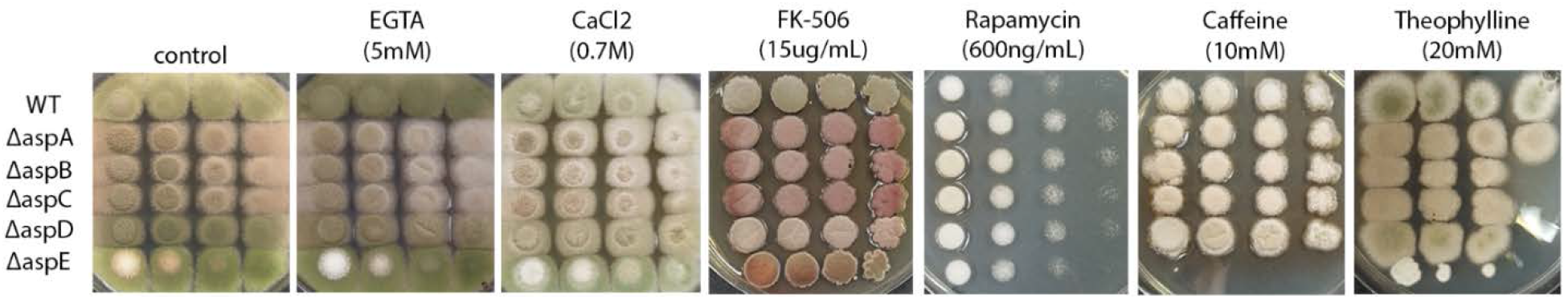
Core septin null mutants are not sensitive to treatments which disrupt the Ca2+/Calcineurin or TOR pathways. AspE is hypersensitive to treatments which disrupt the TOR pathway. WT and septin null mutants Δ*aspA^cdc11^*, Δ*aspB^cdc3^*, Δ*aspC^cdc12^*, Δ*aspD^cdc10^*, and Δ*aspE* were tested for sensitivity by spotting decreasing spore concentrations on complete media plates with or without Calcium/Calcineurin pathway-disturbing agents (EGTA, CaCl2, and FK-506) or Target of Rapamycin (TOR) pathway-disturbing agents (Rapamycin, caffeine, and theophylline). Differences in colony color result from changes in spore production, spore pigment, and production of secondary metabolites under stress. Spore concentrations were [1^7^ conidia/mL – 10^4^ conidia/mL] for all assays in figure. N=3

**S5 Fig.**
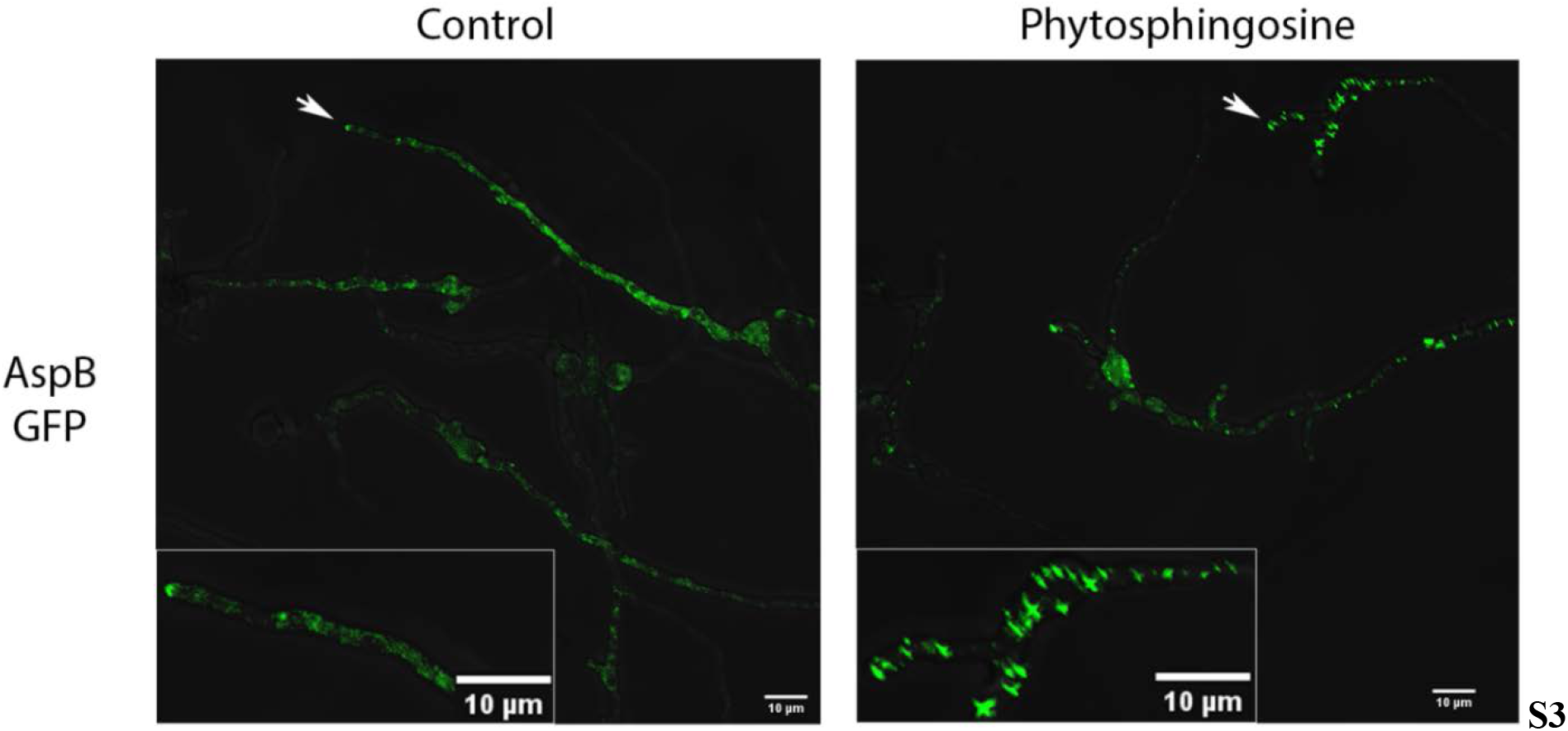
Septin AspB^Cdc3^-GFP localization disrupted by sphingolipid biosynthesis intermediate phytosphingosine. AspB^Cdc3^-GFP strain incubated in liquid media for approximately 16h and imaged 180 minutes after replacing with fresh media containing sphingolipid biosynthesis-inhibiting agents. Representative images are shown from three independent biological replicates, with ≥100 cells observed. (Left Panel) AspB-GFP in vehicle control treatments and (Right Panel) phytosphingosine (15μM) treatment. Enlarged section of micrographs from each picture to better visualize pattern of fluorescence. White arrows denote hyphae which are highlighted in enlarged images. Imaging conducted with Deltavision I deconvolution inverted fluorescence microscope. Scale bars = 10μm. N=3

**S6 Fig.**
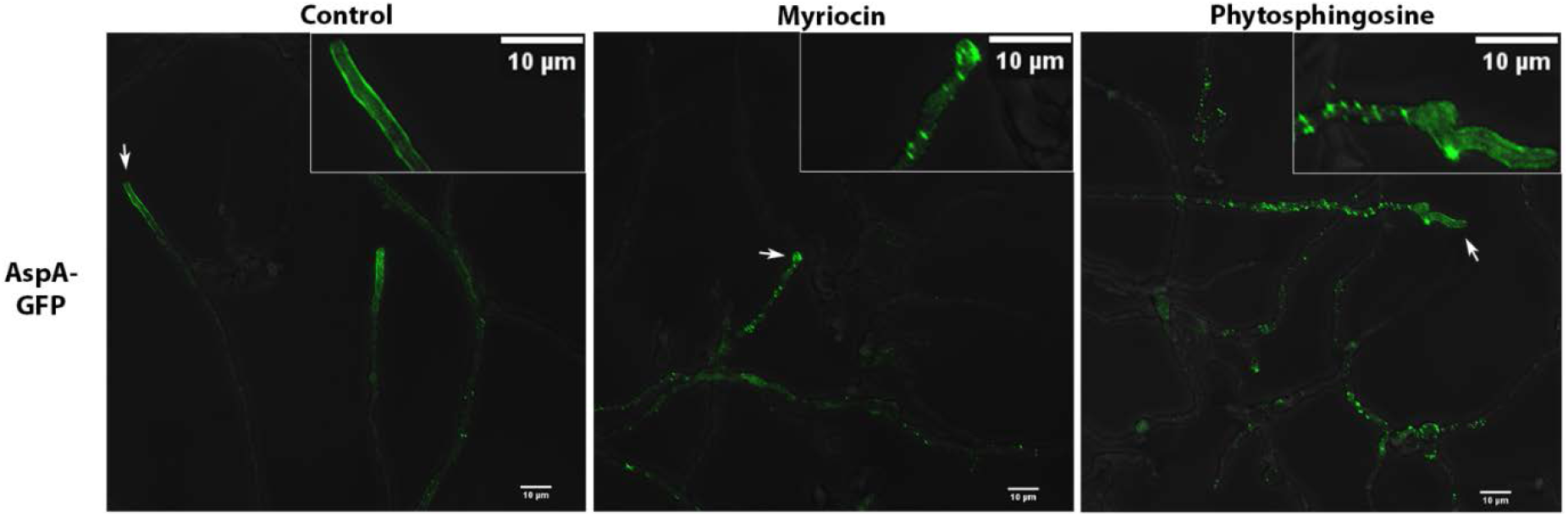
Septin AspA^Cdc11^-GFP localization disrupted by sphingolipid biosynthesis inhibitors. AspA^Cdc11^-GFP strain incubated in liquid media for approximately 16h and imaged 180 minutes after replacing with fresh media containing sphingolipid biosynthesis-inhibiting agents. Representative images are shown from three independent biological replicates, with ≥100 cells observed. (Left Panel) AspA-GFP in vehicle control treatment, (Middle Panel) Myriocin (17.5μg/mL) and (Right Panel) phytosphingosine (15μM) treatment. Enlarged section of micrographs from each picture to better visualize pattern of fluorescence signal. White arrows denote hyphae which are highlighted in enlarged images. Imaging conducted with Deltavision I deconvolution inverted fluorescence microscope. Scale bars = 10μm. N=3

**S1 Table.**
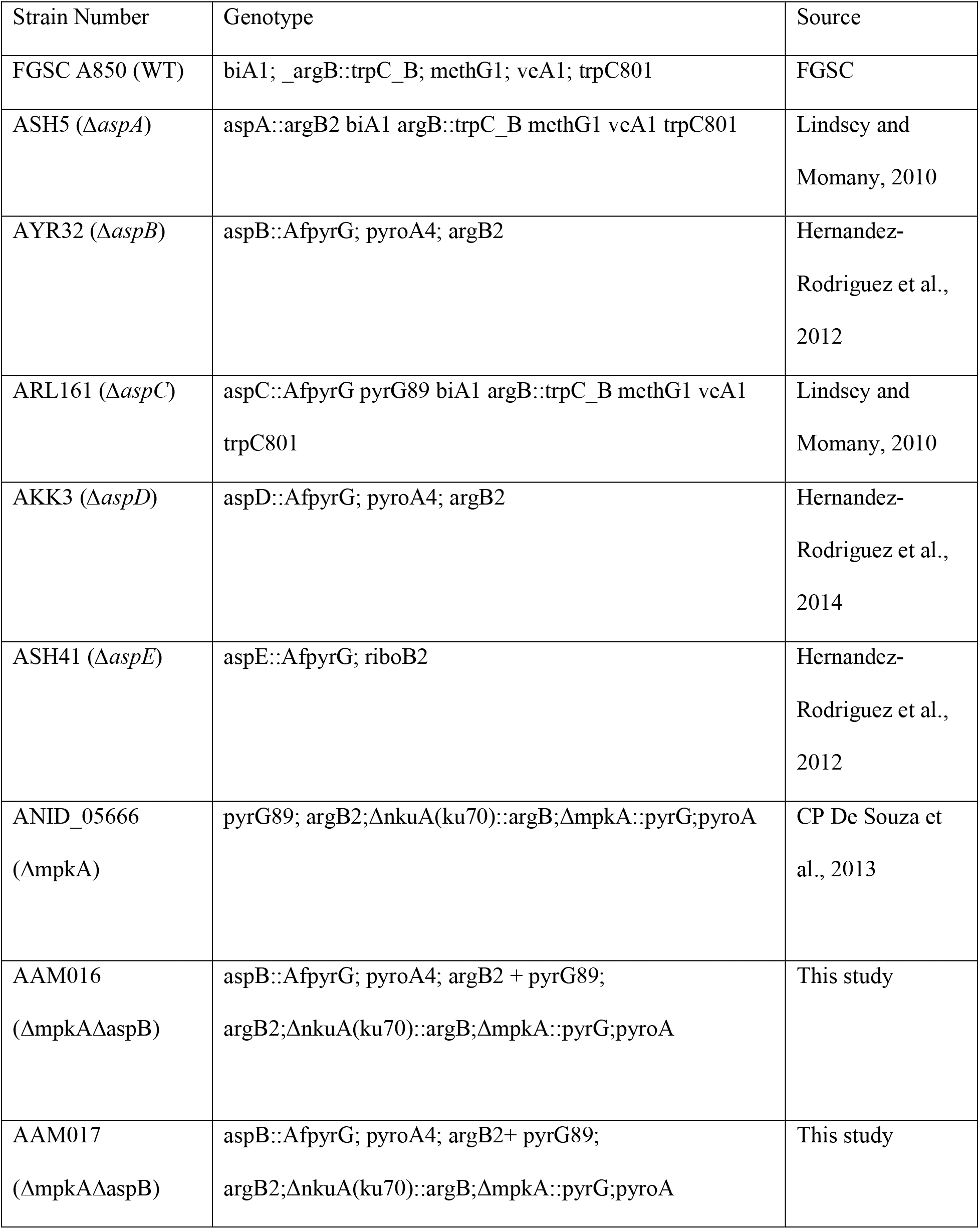
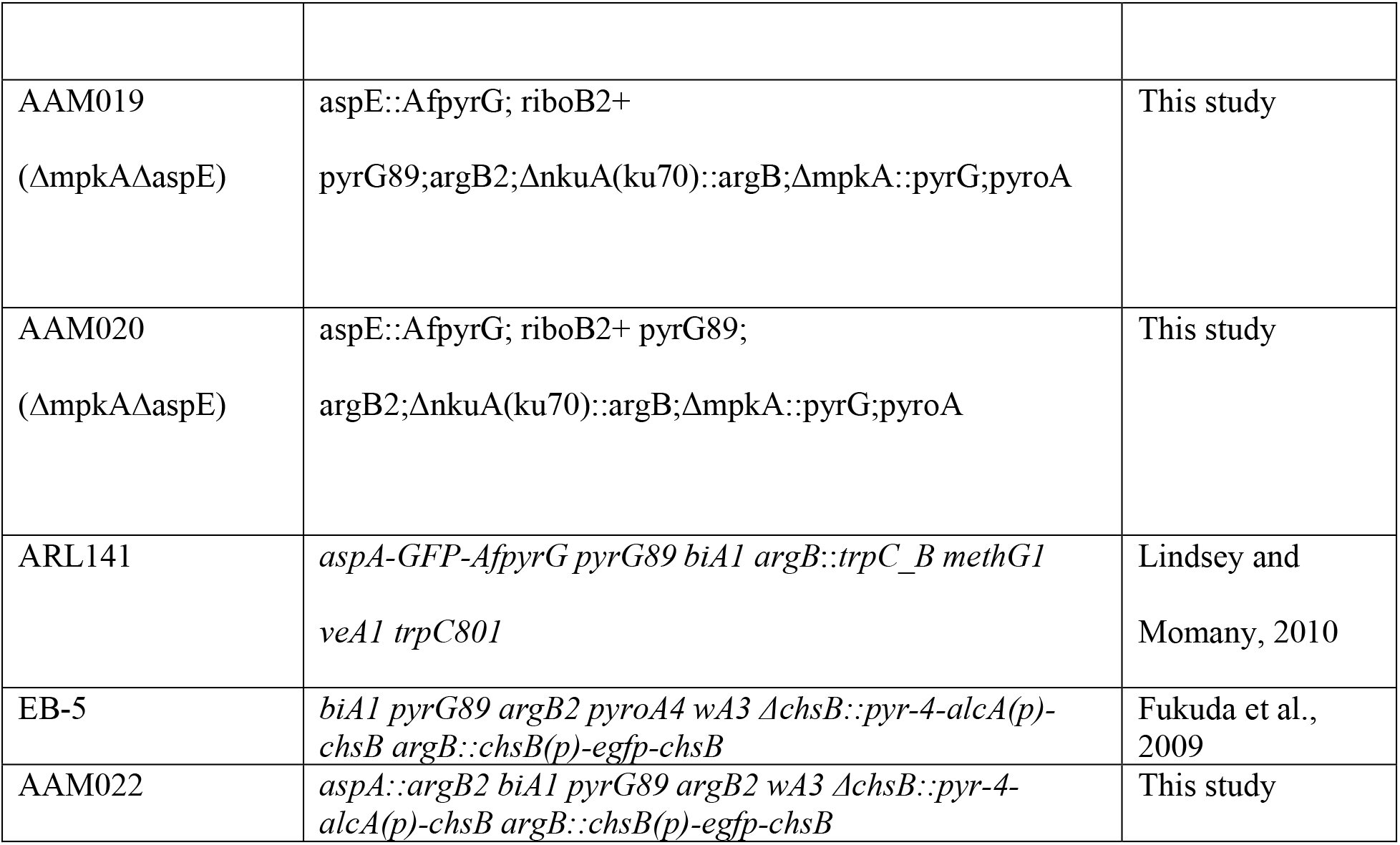
List of fungal strains used in this study.

**S2 Table.**
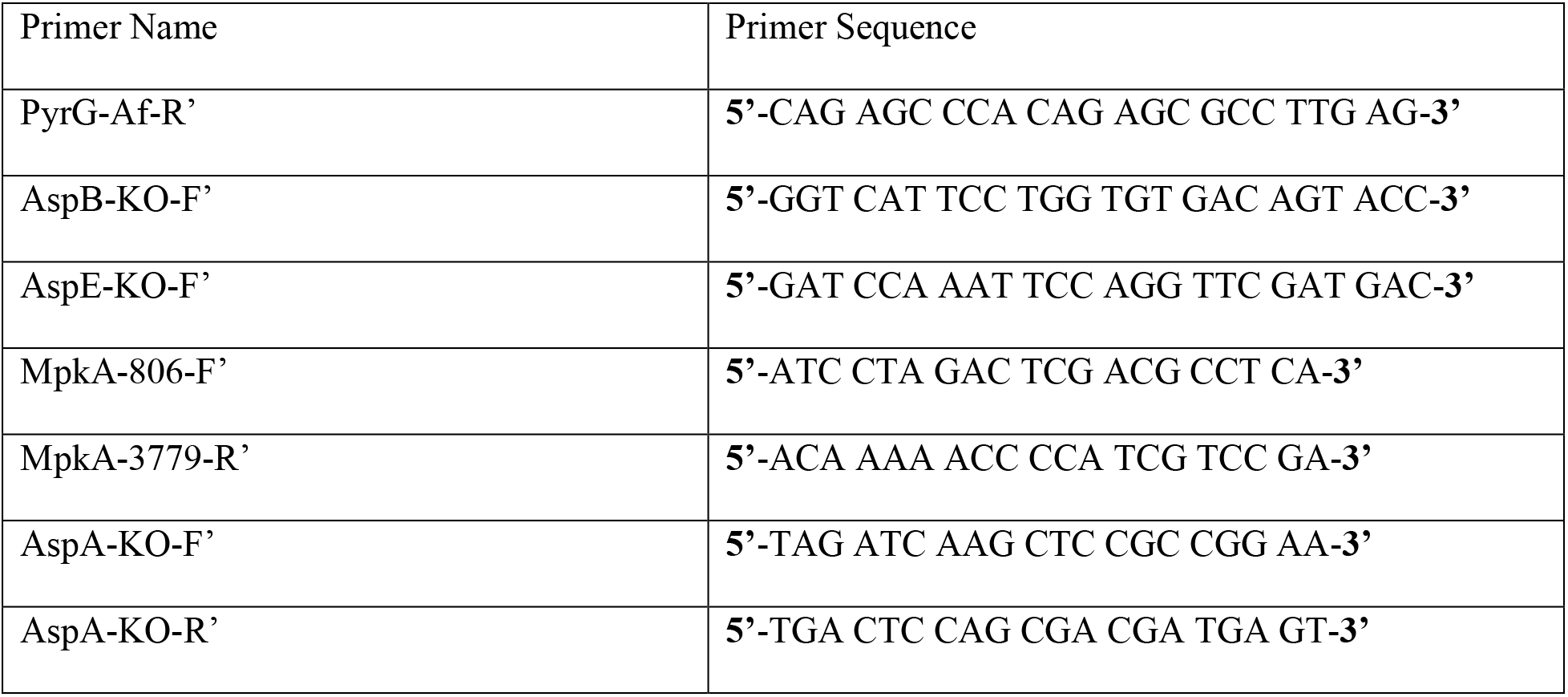
List of primers and sequences used in this study.

